# Spatio-temporal dynamics of the fibrotic niche in cardiac repair

**DOI:** 10.1101/2024.11.10.622609

**Authors:** Andy Shing Fung Chan, Joachim Greiner, Tomas Brennan, Ankit Agrawal, Helene Hemmer, Katrin Sinning, Lisa Marschhäuser, Wing Lam Cheung, Zafar Iqbal, Alexander Klesen, Thomas Seidel, Martin Vaeth, Eva Rog-Zielinska, Peter Kohl, Franziska Schneider-Warme, Dominic Grün

## Abstract

The heart is one of the least regenerative organs in humans, and ischemic heart disease is the leading cause of death worldwide. Understanding the cellular and molecular processes that occur during cardiac wound healing is an essential prerequisite to reducing health burden and improve cardiac function after myocardial tissue damage. By integrating single-cell RNA-sequencing with imaging-based spatial transcriptomics, we reconstructed the spatio-temporal dynamics of the fibrotic niche after ventricular injury in adult mice. Our analysis reveals dynamic regulation of local cell communication niches over time. We identified interactions that regulate cardiac repair, including fibroblast proliferation silencing by *Trem2^high^* macrophages that prevents excessive fibrosis. Moreover, we discovered a rare population of dedifferentiating cardiomyocytes during early post-lesion stages, which was sustained by signals from myeloid and lymphoid cells. Culturing non-regenerative mouse cardiomyocytes or human heart tissue with these niche factors reactivated progenitor gene expression and cell cycle activity. In summary, this spatio-temporal cell type atlas provides valuable insights into the heterocellular interactions that control cardiac repair.

**Highlights:**

- scRNA-seq and in situ sequencing reveal spatio-temporal dynamics of heart repair
- Local heterocellular communication niches coordinate overall wound response
- Fibroblast cell cycle silencing by *Trem2^high^* macrophages suppresses excessive fibrosis
- Cardiomyocyte plasticity is promoted by myeloid and lymphoid cells

## Introduction

Myocardial infarction (MI) is the leading cause of death globally, accounting for 16% of all deaths ^1^. Due to the lack of regenerative capacity of adult hearts, post-MI patients often suffer from impaired cardiac output, resulting in a severe decrease in quality of life. Fewer than 1% of adult human CM can proliferate ^2^. In mouse, neonatal hearts can fully regenerate through cardiomyocyte (CM) dedifferentiation,^3–5^ while adult CM have a drastically reduced capacity for dedifferentiation and proliferation ^2,4^. This transition occurs within about seven days after birth ^4^. Upon MI in 7-day-old mice, CM do not proliferate but develop hypertrophy, characterized by an increased cell size, a shift to pro-lipid metabolism, and induced expression of specific genes including *Nppa, Nppb* and *Xirp2* ^4,6^. Despite the general loss of CM dedifferentiation capacity, signalling ligands such as Nrg1, Oncostatin M, and TWEAK can stimulate partial dedifferentiation and proliferation in adult CM ^7–9^, suggesting the necessity of specific niche determinants for cardiac regeneration. However, these signaling factors are insufficient to prevent chronic fibrosis and/or to promote regeneration in the natural post-MI environment. As the adult heart cannot regenerate, formation of a permanent scar must be tightly regulated to avoid impairment of cardiac function.

Fibrotic scar formation involves a complex, time-dependent communication network of different cell types. Following MI, myeloid cells infiltrate the tissue, secrete proinflammatory cytokines such as IL-6, IL-1β and TNFα ^10^, and clear necrotic myocardium in the damaged region ^11^. Ly6C^high^ CCR2^high^ monocyte-derived macrophages (mo/MP) transition from a pro-inflammatory to a pro-reparative state and activate conversion of fibroblasts (FB) into myofibroblasts (myoFB) ^2^. myoFB migrate into the ischemic zone (IZ) and deposit extracellular matrix (ECM) proteins, forming a fibrotic scar ^12^. While scar tissue is essential for wound closure and mechanical stability,^13^ excessive fibrosis can lead to impaired electrical conduction, reduced ejection function, and heart failure ^14^. Fine-tuning FB activation and ECM deposition is therefore crucial for optimal healing outcomes. Despite some insights having been gained into signalling interactions driving cardiac wound healing ^2^, morbidity and mortality due to adverse left ventricle (LV) remodelling remain high: 20% of patients develop heart failure within 12 months post MI ^15^.

To improve our understanding of the molecular and cellular processes underlying fibrotic scar formation, we have established a single-cell resolution spatio-temporal atlas of post-lesion mouse hearts by integrating single-cell RNA-seq (scRNA-seq) and high-resolution spatial transcriptomics. We infer changes in cell states and tissue architecture during the major stages of wound healing (i.e. early inflammation, scar formation, and maturation). The single-cell spatial resolution reveals the niche composition of the lesion, exposing local cell state dependencies and signalling interactions during scar formation. We report for the first time a dynamic macrophage–fibroblast crosstalk in the late stages of healing that may prevent excessive fibrosis, and we describe the signalling niche of a rare population of dedifferentiated CM in the early to middle stages of healing, suggesting that a remnant regenerative response may persist in the heart between neonates and adults. All data can be interactively explored and visualized in our online resource: https://www.wuesi.medizin.uni-wuerzburg.de/cardiac_spatiotemporal_atlas/

## Results

### A spatio-temporal atlas of cardiac scar formation at single-cell resolution

To explore the spatio-temporal dynamics of cardiac lesion formation at single-cell resolution, we utilized two common mouse models to introduce ventricular lesions: ligation of the left anterior descending coronary artery followed by reperfusion ^16^ (LAD) and cryoablation of the LV ^17^ (Cryo). LAD leads to an ischemic wound in the LV that resembles acute, reperfused MI in the human heart. During Cryo, tissue is frozne with a cryo-probe to induce severe local lesioning of the left LV free wall. Although the Cryo wound does not originate from ischemic damage, its size and location are more consistent across animals compared to the LAD intervention (Fig. 1A).

**Figure 1.**
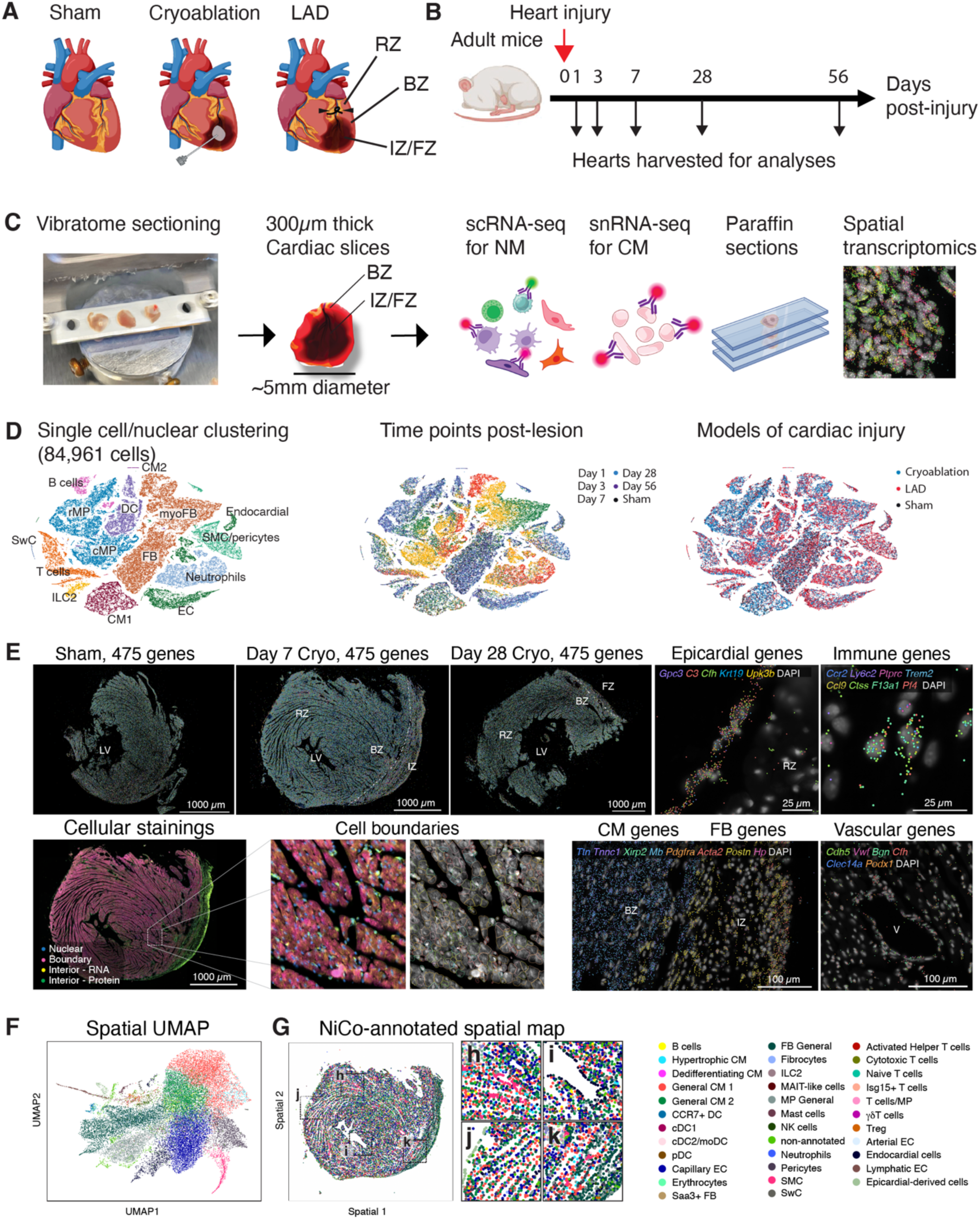
Harvesting of post-lesion cardiac cells for spatio-temporal single-cell transcriptomic characterization. **A – C**, schematic diagrams of surgery models and responses post-lesion (**A**), experimental design (**B**) and heart sample processing procedure (**C**) (Methods). **D**, UMAPs of cells (NM) and nuclei (CM) from all time points and lesion models, composed of 84,961 cells/nuclei. Left, major cell types; center, time points; right, surgery models. **E**, Spatial data from sham day 7 (Sham), Cryo day 7 (Day 7) and day 28 (Day 28) transverse cardiac sections. Top left, all detected transcripts from 475 genes. Right, magnified regions of the day 7 heart, where selected epicardial, vascular, immune, FB and CM genes are shown. Bottom left, cellular counterstaining for cell boundary segmentation. **F and G**, spatial UMAP (**F**) and spatial map (**G**) of day 7 heart highlighting cell type annotations (simplified annotation scheme, Methods) by NiCo algorithm.

After animals had recovered from lesion induction, we isolated and vibratome-cut the LV into tissue slices of 300µm thickness. Slices had a radius of ∼5mm, and were centered on the ischemic zone (IZ, corresponding to days 1 to 7 of the wound healing process) or fibrotic zone (FZ, corresponding to days 28 to 56 of the wound healing process), surrounded by border zone (BZ) tissue (Fig. 1B). For scRNA-seq, CM and non-myocytes (NM) were isolated from cardiac slices obtained at 1, 3, 7, 28 and 56 days post surgery (Fig. 1B,C) by enzymatic digestion, followed by antibody labeling and FACS sorting. After CM exclusion, we combined unbiasedly sorted samples with samples enriched for endothelial cells (EC), smooth muscle cells (SMC), pericytes and Schwann cells (SwC, CD31+/CD146+), dendritic cells (CD45+CD14−CD11c+) and lymphocytes (CD45+CD14−CD11c−) (Fig. S1A, Methods). We also isolated DAPI+PCM-1+ nuclei for single-nucleus RNA-sequencing (snRNA-seq) of CM (Fig. S1B, Methods). Finally, we performed spatial transcriptomics using *in situ* sequencing at single-molecule resolution (10x Xenium) on 5µm sections of paraffin-embedded left ventricles (Methods).

Our single-cell transcriptome atlas contains 84,961 NM cells and CM nuclei from all timepoints (Fig. S1C). Using established marker genes, we annotated major cell populations as CM, FB/myoFB, capillary EC, endocardial EC, SMC/pericytes, neutrophils, circulatory macrophages (cMP), resident macrophages (rMP), B cells, T cells, innate lymphoid cell type 2 (ILC2), and SwC (Fig. 1D and Fig. S1D). Some of these cell types (e.g. MP and FB/myoFB) showed pronounced heterogeneity across different timepoints. However, cells obtained from LAD and Cryo were well mixed in the t-distributed stochastic neighbor embedding (tSNE) but were separated from sham, suggesting that the reparative processes of the two lesion models were generally similar (Fig. 1D). This was supported by the presence of only 22 differentially expressed genes (DEG; *p*-adjusted <0.05) when comparing LAD and Cryo lesions at the pseudobulk level (Fig. S1E). Gene set enrichment analysis revealed condition-enriched pathways related to muscle contraction and antigen processing for Cryo, and respiratory electron transport/ATP synthesis and mRNA splicing for LAD (Fig. S1F). However, hypoxia-related genes such as *Hif1a, Ubb* and *Psm*-gene family genes were upregulated in the two models when compared to sham (Fig. SG-I). These data suggest that both lesion types can represent physiological MI models, characterized by similar molecular remodeling. Nevertheless, slight transcriptomic differences were detected among day 3-7 myoFB, as discussed below.

We selected an early (day 7) and a late (day 28) timepoint of the Cryo model along with a day 7 post-sham-surgery control sample for spatial analysis of 475 genes; genes were selected based on cell type marker genes derived from the sequencing data (Fig. 1E, Methods). For each sample, the field of view covered >14,000 cells (Fig. S1J,K). To validate the reliability of transcript detection and gene decoding, marker genes of epicardial cells, immune cells, FB, CM, and vascular cells were visualized (Fig. 1E). Epicardial genes such as *Gpc3* and *Upk3b* were restricted to the epicardial region. Vascular genes such as *Cdh5* and *Vwf* were localized in vessels, while immune markers (such as *Ccr2, Ly6c2, Trem2* and *F13a1*) and FB genes (such as *Pdgfra, Acta2* and *Postn*) were enriched in the IZ/FZ. CM genes (such as *Ttn, Tnnc1* and *Xirp2*) were restricted to the BZ and remote zone (RZ) (Fig. 1E).

These patterns were consistent with known cell type dynamics following MI, specifically cell death of CM and recruitment of FB and immune cells to the lesion site (Fig. 1A). Cells were segmented by the Xenium pipeline based on fluorescent staining of nuclei, cell membranes, intracellular RNA and proteins (Fig. 1E). Using NiCo ^18^, an algorithm for the integrated analysis of single-cell resolution spatial transcriptomics and matched sc/snRNA-seq reference data, we annotated segmented cells in the spatial modality for downstream analyses of niche interactions (Fig. 1F,G and Fig. S2).

### Dynamic waves of immune cell populations during scar formation

The combination of unbiased sampling and enrichment of rare cell populations yielded a comprehensive spatio-temporal atlas of immune cell type dynamics during scar formation (Fig. 2A-C and Fig. S1A). Most myeloid cell types, including neutrophils, MP, dendritic cells (DC) and mast cells, were highly enriched on days 1 – 7; this was followed by expansion of lymphoid populations on day 7, and restoration of sham-like cell type proportions on days 28 and 56 (Fig. 2C and Fig. S3A). Consistent with limited global expression differences between Cryo and LAD (Fig. 1G), cell type proportions were also similar for the two lesion models, yet different from sham (Fig. S3B). NiCo recovered all major immune cell populations in the spatial modality (Fig. 2D and Fig. S3C – E). The majority of immune cells, dominated by MP, were localized to the IZ on day 7, whereas the immune cell density was globally reduced on day 28 and even lower in sham (Fig. 2D and Fig. S3F,G).

**Figure 2.**
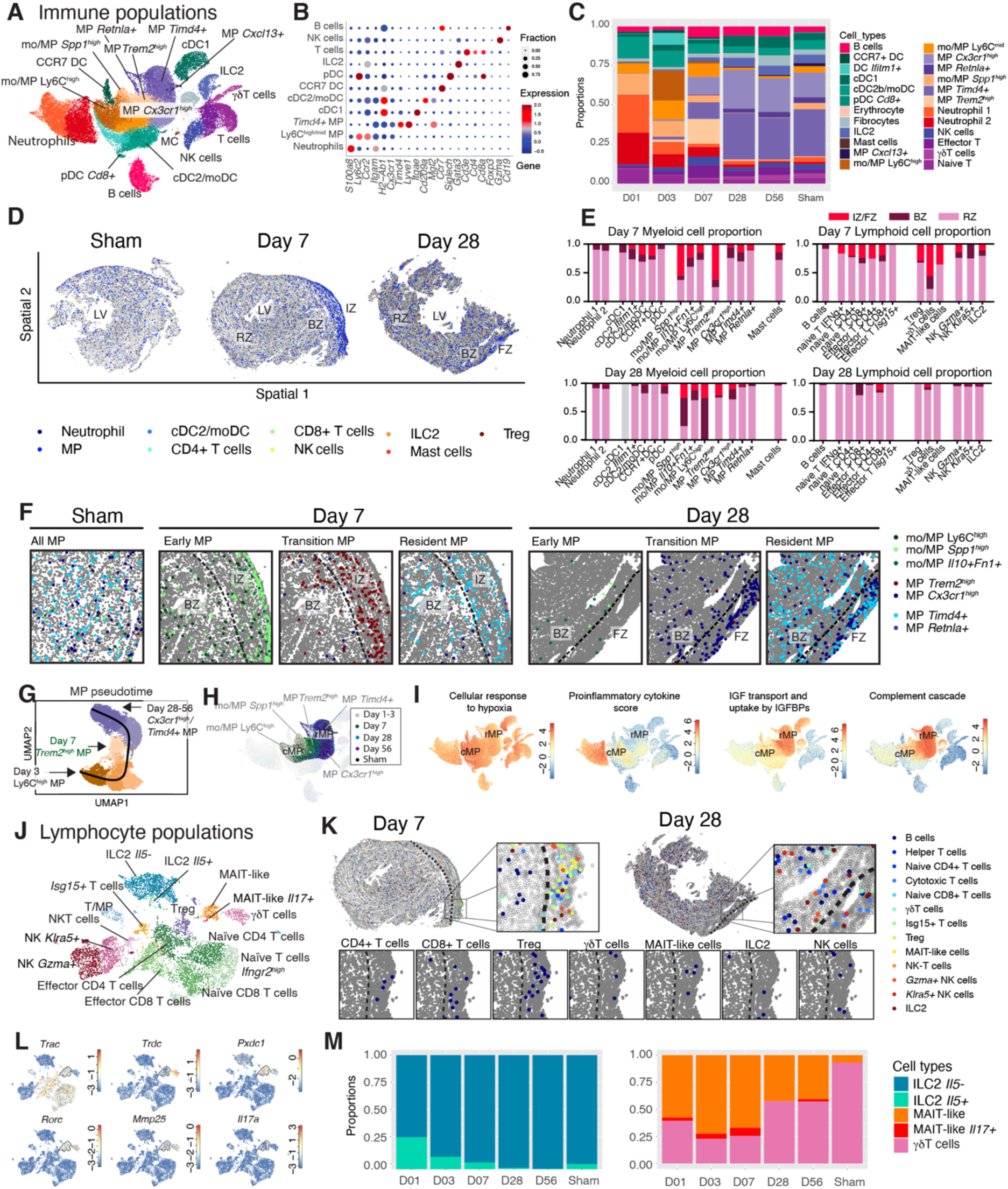
Spatio-temporal cell type dynamics of the immune compartment. **A**, UMAP of scRNA-seq data of immune populations from day 1 to 56 post lesion. **B**, DEGs for each immune population. Dot size indicates fraction of cells expressing the gene. **C**, immune cell type proportions over time (day 1 to 56) and for Sham. **D – E**, Spatial maps of immune cells for Sham, day 7, and day 28 under the detailed annotation scheme (**D**), with quantification of myeloid/lymphoid cell frequencies across injury or fibrotic (IZ/FZ), border (BZ) and remote zone (RZ) for day 7 and 28 samples (**E**). **F**, spatial distribution of different mo/MP subtypes in the wound regions (or the edge of LV for Sham). Dotted lines separate IZ/FZ and BZ. **G and H**, inferred MP cell fate transition (**G**) and changes in enriched cell states across different time points (**H**). **I**, summed expression of pathway genes enriched in cMP and rMP. **J**, scRNA-seq UMAP of lymphocyte populations that were subclustered from (**A**). **K**, spatial distribution of lymphocyte populations. **L**, UMAP showing gene expression of marker genes of γδT and MAIT cells, and *ll17a*. Highlighted in dotted line: MAIT-like cells. **M**, temporal cell type proportion change of *Il5+* and *Il5-* ILC2 (left) and *Il17-*/*Il17+* MAIT-like cells and γδT cells (right).

### Temporal phenotypic conversion of macrophages post lesion

Throughout the process of scar formation, the turnover dynamics of infiltrating MP from the circulation (cMP) and cardiac resident MP (rMP) as well as their phenotypic modulation are critical to determining the progression and outcome of wound healing. Consistent with previous findings ^10,19,20^, our spatial data of the Cryo heart showed that on day 7 the site of lesion was dominated by *Spp1+* monocyte/macrophages (mo/MP) and *Trem2^high^* MP, while *Cx3cr1^high^* MP became more prevalent on day 28 (Fig. 2E,F). Pseudotime analysis (Methods) revealed a gradual lineage transition of day 1-3 lesion-enriched Ly6C^high^ and *Spp1^high^* cMP states, towards MHCII^high^*Cx3cr1^high^* and *Timd4^high^* rMP phenotypes (Fig. 2G), which were highly enriched on day 28 and 56 and in sham (Fig. 2H). A day 7-enriched *Trem2^high^* population may represent a pivotal transitory population during the conversion from cMP to rMP phenotypes (Fig. 2G).

Differential gene expression analysis (Methods) revealed pronounced differences between rMP and cMP states. cMPs upregulated proinflammatory cytokines (*Il6*, *Il1b,* and *Osm*), glucose metabolism (*Gapdh*, *Pgk1*, *Pgam1,* and *Aldoa*), reactive oxygen species (ROS) detoxification, hypoxic genes and nucleotide di/triphosphates interconversion. rMP were enriched in degranulation-related genes (*Pf4* and *Selenop*), insulin-growth factor (IGF) transport/uptake genes (*IgQp4*, *Igf1* and *Gas6*), and complement cascade genes (*C4b*, *C1qb*, *C1qc* and *Cd81*) (Fig. 2I, Fig. S3H-J). Hence, the observed phenotypic conversion around day 7 silences inflammatory programs and cell stress related to hypoxia and ROS, while pro-repair factors such IGF pathway genes are enhanced.

Moreover, the cMP population undergoes active proliferation after tissue infiltration (days 1– 3), which is suppressed by day 7. In contrast, proliferation of rMP does not change over time (Fig. S3K). The lower cell numbers of the cMP at mid-to-late stages of lesion repair could be attributed to cell cycle suppression and phenotypic conversion into rMP, whereas the expansion of the rMP population most likely results from phenotypic conversion of cMP.

### Dynamic modulation of cytokine production by innate lymphocytes and non-conventional T cells

Our dataset revealed pronounced time-dependent heterogeneity of lymphocyte populations comprising conventional and unconventional T cells, NK cells and ILC2 (Fig. 2J-L and Fig. S3D, S4A-D and F). Except for naïve T cells, most lymphocyte populations expanded and peaked between day 7 and 28 (Fig. S4B,D), were spatially-enriched in the IZ on day 7, and were cleared by day 28 (Fig. 2E,K). For ILC2, we discriminated a minor but early (day 1) wound-responding *Il5+* population from *Il5−* subtypes. (Fig. 2M). While T cell-regulatory (*Cd274* and *Ctla4*) and RHO GTPase cycle genes (*Akap13* and *Wipf1*) were upregulated in *Il5-* ILC2s, the *Il5+* subtype was more specialized in ROS sensing, RUNX2 regulation, MHCII antigen presentation, and cell stress response (Fig. S4E), suggesting a temporal shift from immune activation to suppressive functions.

We recovered populations of γδT cells and mucosal-associated invariant T (MAIT)-like cells. While both cell types expressed known MAIT and γδT cell markers such as *Rorc*, *Pxdc1,* and *Mmp25*, only γδT cells expressed γ/δ TCR subunit genes such as *Tcrg-C1*, *Trdc,* and *Trdv4*, and the scavenger receptor cysteine-rich family gene *Cd163l1*. MAIT-like cells expressed α/β TCR genes such as *Trac* and *Scart2*, upregulated OXPHOS, IL-2, and IL-7 pathways, and were the only lymphocytes in our dataset expressing high levels of *Il17a* (Fig. 2L and Fig. S4F-H), a crucial proinflammatory mediator for neutrophil recruitment and T cell activation. Expansion of the MAIT-like population was observed during earlier stages (days 1–7), followed by expansion of

γδT cells (days 7–56) (Fig. 2M).

### Spatio-temporal dynamics and phenotypic conversion of fibroblasts post lesioning

FB are one of the major responders to lesion induction. Our single-cell sequencing atlas revealed a heterogeneous cell state composition, including the presence of several small clusters such as monocyte-derived fibrocytes ^21,22^ and a *Saa3+* (*Ptprc*−) FB population (Fig. 3A-B). The main clusters can be collectively grouped into quiescent FB, early *Ccl2+* FB, myofibroblasts (myoFB) and transition FB (tFB). Quiescent FB were dominant in sham, day 28 and 56 hearts, while *Ccl2+* FB, myoFB and tFB were more abundant on days 1, 3 and 7, respectively (Fig. 3C).

**Figure 3.**
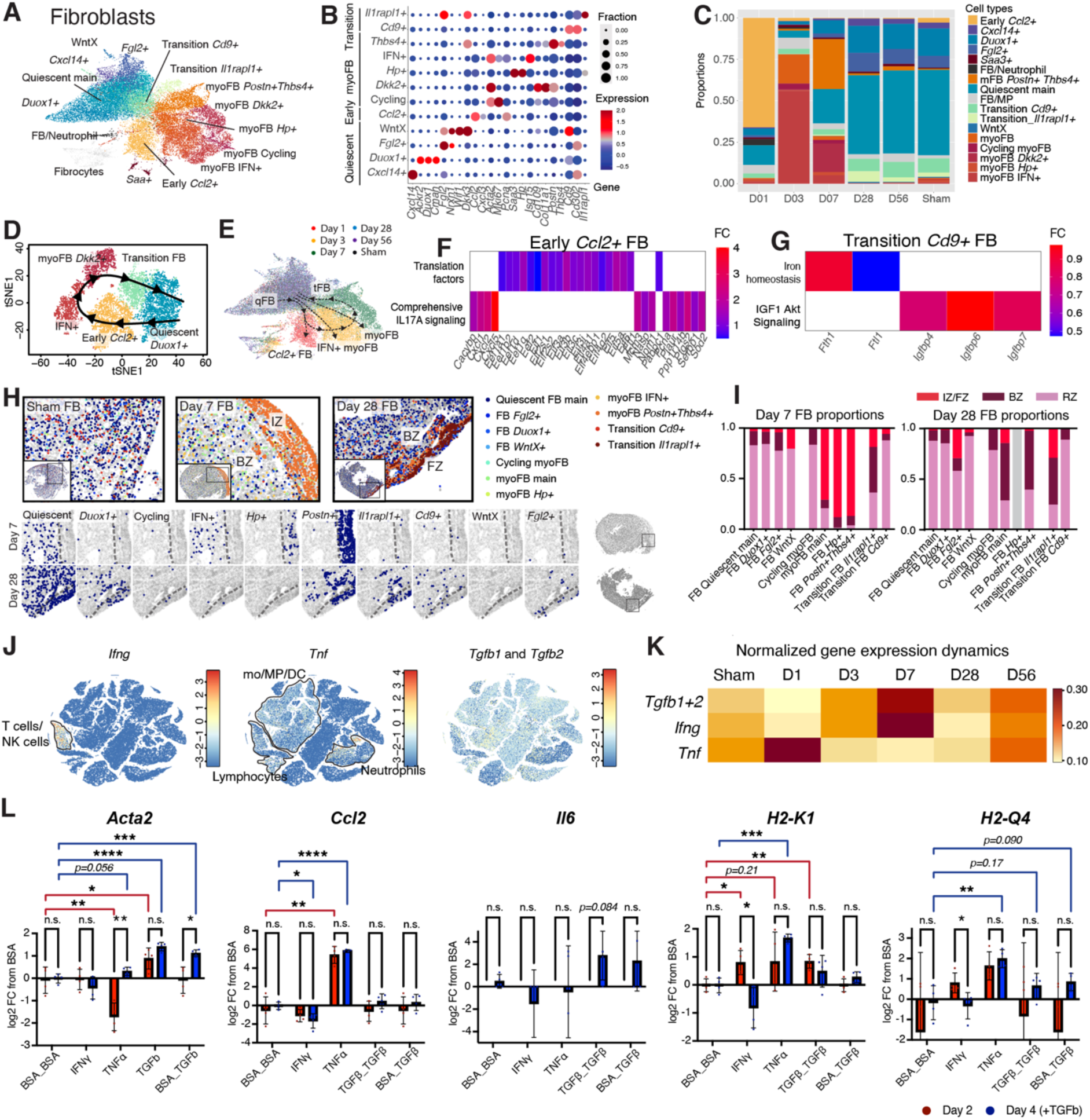
Spatio-temporal dynamics and phenotypic conversion of cardiac fibroblasts. **A**, UMAP of scRNA-seq data for FB subtypes from day 1 to 56 post-lesion. **B**, DEGs for each FB subtype, grouped into “Quiescent”, “Early”, “myoFB” and “Transition” states. Dot size indicates fraction of cells expressing the gene. **C**, cell type proportion of FB subtypes across days 1 – 56 and Sham. **D**, pseudotime FB cell fate transition during repair. **E**, enriched cell states over time. “qFB” Quiescent, “*Ccl2*+” Early *Ccl2*+, “tFB” Transition FB. **F**, heatmap of pathway enrichment analysis in *Ccl2+* (**F**) and Transition *Cd9+* FB (**G**) (ReactomePA, DEG log2-foldchange > 0, *p* < 0.05). **I**, annotated spatial map showing distribution of FB subtypes in day 7 and 28 hearts. **J**, tSNE of scRNA-seq data for all cells showing expression of IFNγ, TNFα and TGFβ genes. **K**, heatmap of normalized gene expression of the genes in (**J**) across time. **L**, qRT-PCR of day 2 and 4 cultured FB for FB cell state markers *Acta2* (myoFB), *Ccl2* (*Ccl2+* FB), *Il6* (*Ccl2+* FB and myoFB), *H2-K1* and *H2-Q4* (IFN+ MHCII^high^ FB). Statistical significance was calculated by unpaired two-tailed t-test between two groups. * *p* < 0.05, ** *p* < 0.01, *** *p* < 0.001, and **** *p* < 0.0001; ns, not significant.

Minor transcriptomic differences were detected between myoFB responding to Cryo and LAD, such as upregulation of genes related to AP-1, MAPK kinases, and IL-17 signaling pathways in LAD-responding myoFB (Fig. S5A-C). However, no major differences were found between post-LAD and post-Cryo. Quiescent FB could be further subclassified into multiple transcriptomically different states, such as WntX ^23^, *Fgl2+*, *Cxcl14+* and *Duox1+* populations. Upregulation of proinflammatory cytokines and chemokines such as *Ccl2, Ccl7* and *Il6* was observed in early activated *Ccl2+* FB ^24^, which was partially shared with myoFB (Fig. S5D). *Acta2+* myoFB were highly abundant from days 3 to 7, comprising interferon (IFN)-responsive (IFN+), cycling (day 3-enriched), *Hp+*, *Dkk2+,* and *Postn+Thbs4+* (day-7-enriched) ^25,26^ subtypes, with gradual increase in the expression of collagen genes such as *Col1a1*. The IFN+ subtype, in particular, showed higher expression of MHCII genes for presenting antigens to CD4+ T cells ^27^ (Fig. S5D).

Previous studies have shown that *Ccl2+* (proinflammatory), IFN+ (MHCII^high^) and Collagen^high^ FB/myoFB states can be individually activated from the quiescent state by TNFα, IFNγ, and TGFβ, respectively ^24,27,28^. In our data, TNFα is highly abundant only on day 1 post lesion, majorly expressed by mo/MP, followed by increasing levels of IFNγ (T/NK cells) and TGFβ (Fig. 3J,K), aligning with the change of FB subtype abundance. Slingshot ^29^ pseudotime analysis reconstructed a trajectory connecting quiescent to early *Ccl2+* FB on day 1, followed by myoFB on days 3 and 7, and further transitioning via *Cd9+* tFB back to quiescent-like *Duox1+* FB (Fig. 3D,E), suggesting that these states may align along a continuous trajectory. *Ccl2+* and *Cd9+*/*Il1rapl1+* tFB are the two intermediates that bridge quiescent and activated states (from quiescent to myoFB, and vice versa, respectively). *Ccl2+* FB promoted protein translation and IL17A signaling responses, suggesting an inflammation-driven, growth-active machinery. Whereas *Cd9+* FB upregulated iron homeostasis, a signature of cellular senescence ^30^, and IGF1 responses, which may suppress fibroblast proliferation ^31^ (Fig. 3F,G). This suggests that different molecular mechanisms govern the two conversion processes.

Except for *Ccl2+* FB, all FB/myoFB states were also observed in the spatial data (Fig. S5E,F). Quiescent FB were only found in the BZ and RZ on day 7 post lesion (Fig. 3H). myoFB were detected both inside and outside of the lesioned region. While IFN+ myoFB were restricted to the BZ, *Hp+* and *Postn+Thbs4+* states were exclusively localized to the IZ (Fig. 3H,I) suggesting a microenvironment-dependent differentiation of FB along with their migration towards the IZ. On day 28, both of the above-mentioned myoFB populations were depleted from the FZ and were replaced by high numbers of *Il1rapl1+*/*Cd9+* tFB and quiescent FB, indicating a transition towards a de-activated state (Fig. 3H).

We next sought to functionally validate the plasticity of cardiac FB suggested by the pseudotime analysis in response to stimuli dominant in the post-lesion heart (Fig. 3J,K). FB isolated from adult mouse hearts were first exposed to either IFNγ, TNFα or TGFβ alone, followed by secondary TGFβ stimulation (Fig. S5G). Notably, all FB populations express receptors of all these cytokines (Fig. S5H). Exposure to TGFβ resulted in upregulation of the myoFB marker *Acta2*, indicating direct differentiation of FB to myoFB (Fig. 3L). These myoFB also expressed low levels of *Il6* and MHCII genes *H2-K1* and *H2-Q4*, aligning with the single-cell data. Exposure to IFNγ and TNFα induced IFN+ MHCII^high^ and *Ccl2+* FB phenotypes, respectively. Note that TNFα alone was sufficient to induce both phenotypes, but not IFNγ. Secondary stimulation of these IFNγ/TNFα-exposed FB with TGFβ resulted in downregulation of MHCII genes in IFN+ FB, and upregulation of *Acta2* in *Ccl2+* FB (Fig. 3L). In summary, these data indicate that (quiescent) FB can directly differentiate into *Ccl2+* FB, IFN+ FB, and myoFB, and that *Ccl2+* FB can further differentiate into myoFB in response to TGFβ.

### Vascular cells and Schwann cells mount an IFN response at early stages post lesion

In addition to the FB response, vascular cells and SwC also play key roles in post-lesion remodeling. Vascular cells can be generally grouped into EC, pericytes, and SMC. EC can be further subclassified into arterial^26^, capillary, endocardial, lymphatic and *Areg*+ states (Fig. 4A-C, and Fig. S6A-D). Similar to FB, both capillary EC and quiescent pericytes were activated upon injury to proliferate on days 1 – 3, and acquired an antigen-presenting angiogenic/IFN+ state (*Isg15+ Ifit3+* MHCII^high^) (Fig. 4D-E and Fig. S6B,C). Upon activation, IFN+ EC mount a hypoxic and ROS response, as well as *Runx2* regulation, but show reduced gene expression related to RHO GTPase (Fig. S6E). IFN+ EC were not observed after day 7 (Fig. 4C and Fig. S6B,C). Given their emergence preceding IFNγ signaling (which peaked on day 7; Fig. 3K), IFN+ EC are probably induced by type I IFN signaling. Spatially, IFN+ EC/pericytes were abundantly found in the BZ and RZ on day 7, but were less frequent in the IZ. Instead, day 28 EC mostly comprise quiescent phenotypes in all regions (Fig. 4F-H, Fig. S6A,B,D). An overall increase of EC density on day 28 indicated global angiogenesis. Endocardial cells were mostly restricted to the endocardium lining the LV, with some EC in IZ/FZ also acquiring endocardial-like expression patterns (Fig. 4F,G). Lymphatic EC were already abundant in the IZ and FZ on day 7, and exhibited increased density on day 28, suggesting prolonged lymphoangiogenesis that establishes and maintains lymphatic vessels in the scar region (Fig. 4F,G). Expansion of pericytes and FB populations could be partially fueled by a small population of epicardial-derived (*Msln+ Mgp+*) cells, clustered between IFN+ and FB-like pericytes (Fig. 4A), which may also acquire FB identities via an epicardial-FB/pericyte trajectory (Fig. 4E). Despite appearing in low cell numbers, epicardial cells and their derivatives were found to be enriched not only in the epicardial lining but also within the IZ/FZ, suggesting their active response and recruitment towards the lesion (Fig. 4F,H).

**Figure 4.**
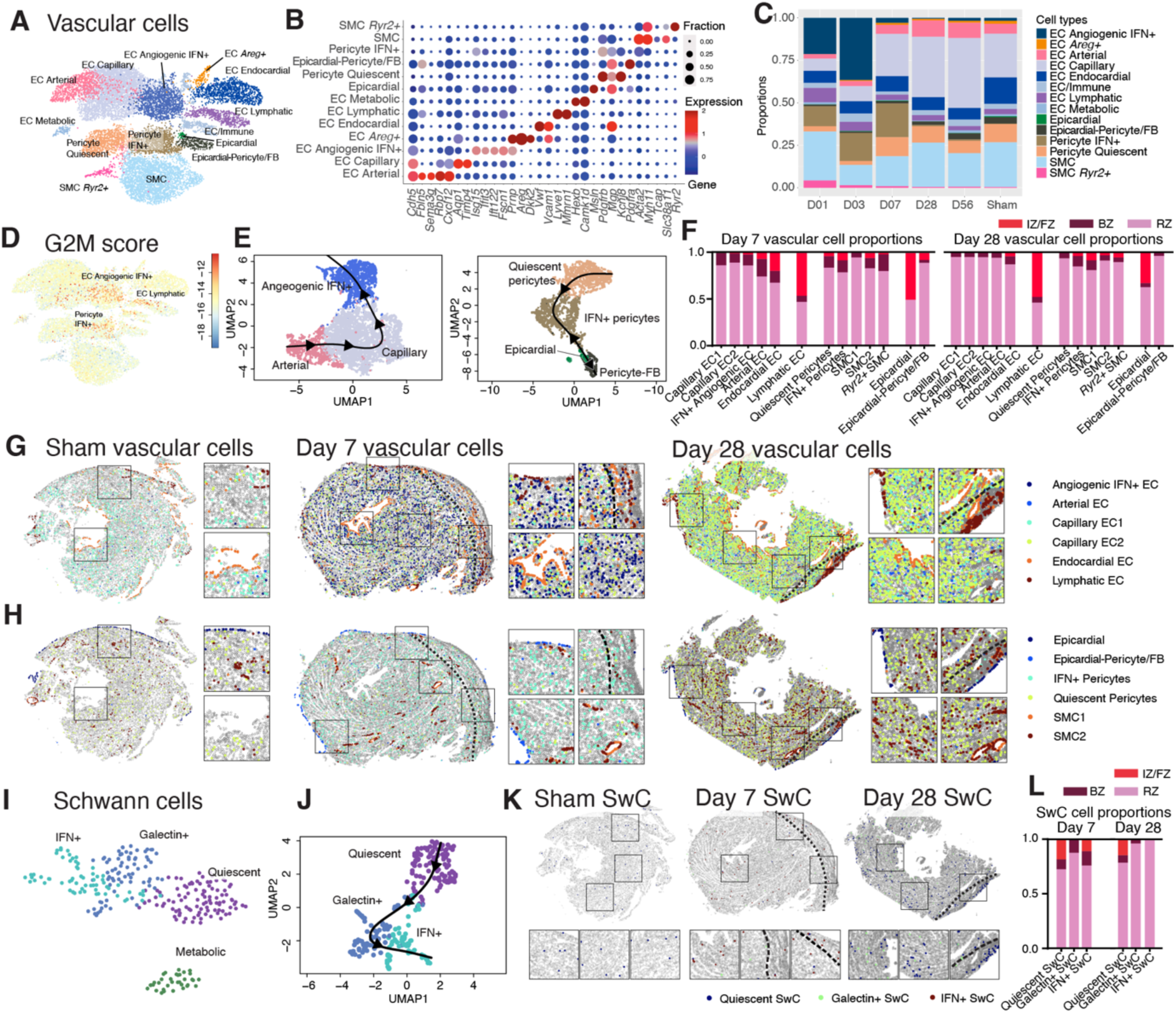
Spatio-temporal dynamics of vascular cells and Schwann cells. **A**, UMAP of scRNA-seq data for vascular populations from day 1 to 56 post-lesion. **B**, DEGs for each vascular subtype. Dot size indicates fraction of cells expressing the gene. **C**, cell type proportion of vascular subtypes across day 1 – 56 and Sham. **D**, G2M score in vascular populations. **E**, pseudotime analyses of EC (left) and epicardial – pericytes/FB (right). **F**, spatial distribution of vascular subtypes on day 7 and 28. **G – H**, spatial maps of EC subtypes (**G**), and epicardial cells, pericytes and SMC (**H**) for day 7 and 28 hearts. **I**, UMAP of scRNA-seq data for SwC subtypes from day 1 to 56 post-lesion. **J**, pseudotime analysis of SwC. **K**, Annotated spatial maps of SwC subtypes. **L**, spatial distributions of SwC subtypes in day 7 and 28 hearts.

Despite being a rare neural cell type in the left ventricle, SwC displayed significant temporal and spatial heterogeneity post lesioning (Fig. 4I,J and Fig. S6H-L). In sham conditions, only a single quiescent SwC state was detected (Fig. S6L,M). Upon lesion induction, SwC became activated into a Galectin+ state (expressing *Lgals3, Cd63,* and *Mt2*) on day 1, transitioning into an IFN+ state in the IZ by day 3; this was similar to the IFN response seen in EC and pericytes (Fig. 4I-L). These states gradually diminished after day 7, replaced by a quiescent state by day 28, aligning with the overall trend of late-stage healing, where most cell types restored pre-wounding phenotypes. Notably, the Galectin+ state expresses a different spectrum of ECM than the quiescent state, such as reduced expression of proteoglycans, laminins, and collagens, but increased fibronectin, tenascin, and fibulin, indicating their role in ECM remodeling (Fig. S6N). The transient activation of SwC and their spatial distribution across BZ and IZ suggest that, alongside vascular cells, they may contribute to the IFN response and wound healing dynamics during post-lesion remodeling.

### Substantial rewiring of the niche architecture during scar formation

The cell state of individual cell types can be substantially affected by heterocellular interactions within their local neighborhoods. We analyzed the spatial data with NiCo ^18^ to investigate significant cell type interaction patterns. NiCo identifies significant niche interactions by training a logistic regression classifier to predict cell type identity from the frequency enrichments of all cell types within their niche across all instances of a cell type. Positive regression coefficients indicate preferential interactions, and NiCo derives a global cell type interaction map from these coefficients for sham, day 7, and day 28 (Fig. 5A-C). Distinct spatial interaction domains were observed on day 7. A “fibrotic niche” was dominated by *Postn+Thbs4+* and *Hp+* myoFB subsets, colocalizing with *Spp1^high^* and *Trem2^high^* MP (as observed previously ^32^) and T lymphocytes such as effector CD4+ T cells, γδT cells and Treg (Fig. 5D). These lymphocytes colocalized with NK cells, ILC2, MAIT-like cells, and DC (including cDC2/moDC and CCR7+ subtypes), forming an antigen-presenting hub (Fig. 5B). In the BZ, abundant hypertrophic CM colocalized with vascular cells and SwC (Fig. 5E). Other CM (such as homeostatic and dedifferentiating CM) formed an interaction hub with capillary EC, naïve T cells, B cells, and DC (including pDC and cDC1 subtypes) in the RZ (Fig. 5F-G). At the outer surface of the heart, epicardial cells co-clustered with epicardial derived pericytes/FB, mast cells, lymphatic vessels and *Spp1^high^* mo/MP (Fig. 5H). The segregation between the fibrotic and the epicardial niche, and between hypertrophic and homeostatic CM niches, were less clear on day 28.

**Figure 5.**
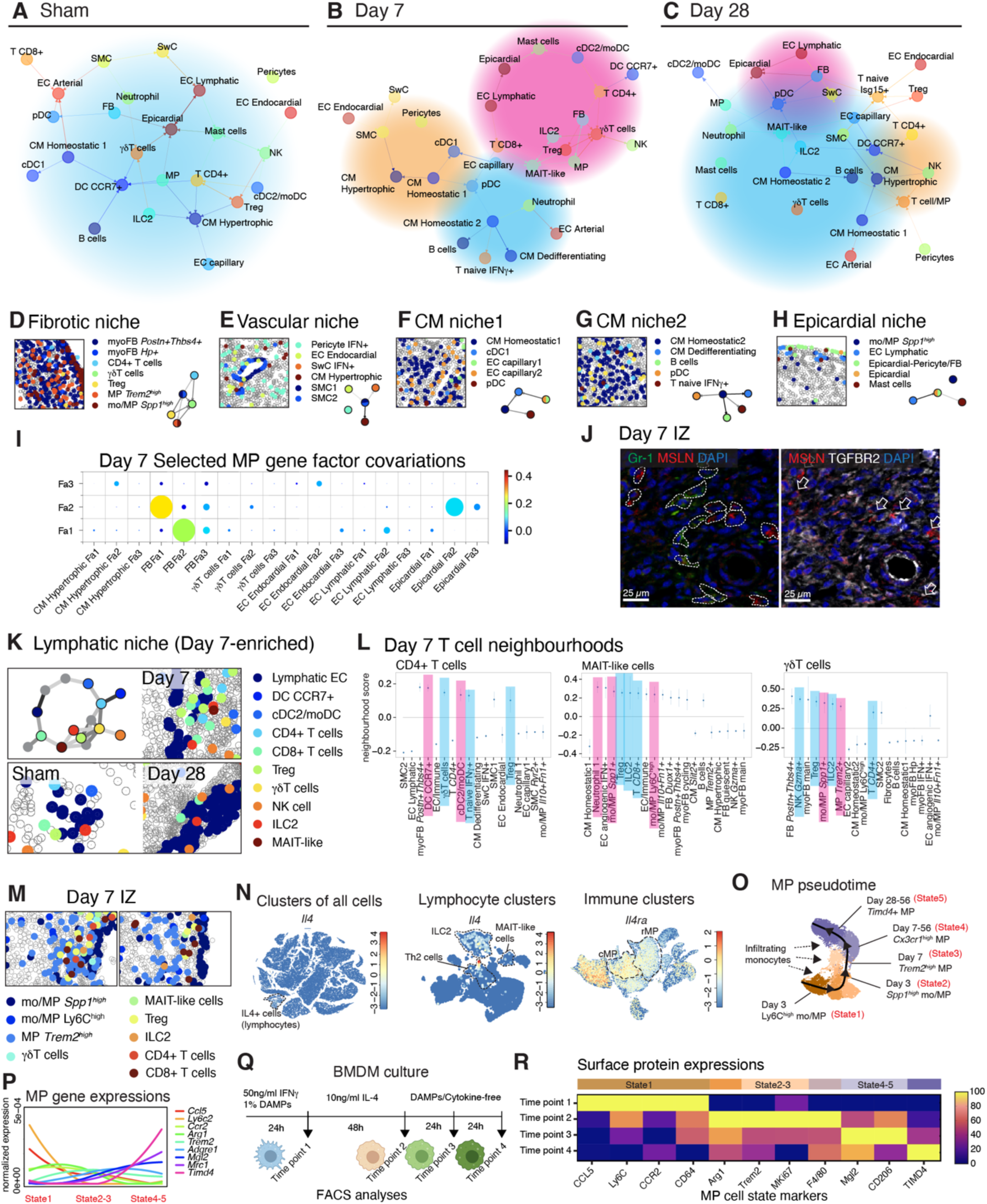
Spatial cell type interactions within immune-vascular niches. **A - C**, NiCo spatial interaction network predictions for Sham (**A**), day 7 (**B**) and day 28 (**C**) hearts. Thickness of edges indicates interaction strength. Arrowheads point from niche cell to central cell. Red shading, IZ/FZ-enriched; orange shading, BZ-enriched; blue shading; RZ-enriched. **D – H**, representative spatial map showing spatial niches in the Day 7 heart, identified in (**B**). **I**, Gene factor covariation dot plot of day 7 MP with other cell types. “Fa”, gene factors. Circle size scales linearly with −log_10_(p-value), and circle color indicates ridge regression coefficients (Methods). **J**, representative IF images of Day 7 heart within IZ, showing colocalization of epicardial-derived cells (MSLN+TGFBR2+) with MP and neutrophils (Gr-1+). White dotted lines highlight cell boundaries (left). Arrows points to epicardial-cells (right). **K**, day 7 lymphatic niche. Upper left: interaction network across lymphocytes and DCs. Others: distribution of these cell types within the niche. **L**, day 7 T cell neighborhoods highlighting myeloid (red) and lymphoid (blue) cells. **M**, representative Day 7 MP-lymphocyte niches. **N**, UMAPs showing *Il4* expression in all cells (left) and lymphocyte populations (middle), and *Il4ra* expression among immune populations (right). **O**, pseudotime cell fate prediction of MP. The lines and arrows indicate potential paths and directions of cell fates. **P**, expressions of cell surface marker genes associated to different MP states. The corresponding surface proteins were quantified in (**Q**). **Q**, schematic diagram of BMDM culture mimicking the time-dependent microenvironmental changes that are exposed to infiltrating MPs (Methods). **R**, FACS analysis of BMDM culture, showing MP state-associated protein expressions.

### Macrophages may promote epithelial-to-mesenchymal transition in epicardial-derived cells

*Ly6C^high^* and *Spp1^high^* mo/MP are recruited mainly on days 1 to 3 after cardiac injury and remain abundant until day 7 (Fig. 2C,F). These MP interact with lymphocytes and multiple non-immune cell types, such as myoFB, EC, hypertrophic CM, EC, pericytes, SMC, Galectin+ SwC, and epicardial cells (Fig. S7A). Beyond preferential niche co-localization, NiCo also predicts covariation of cell states between co-localized cell types. Covariation coefficients are inferred by a linear regression of cell type-specific latent variables, or factors, of a central cell type on latent factors of its neighbors. These latent factors capture cell type-specific transcriptome variability and can be correlated with gene expression across cells to interrogate factor-associated genes for functional enrichment (Methods). Hence, regression coefficients are informative regarding the covariation of particular gene programs in co-localized cells. Applied to MP, NiCo inferred covariation of MP factor (Fa) 2, with Fa2 and 3 of epicardial-derived cells (Fig. 5I). A similar interaction occurs during zebrafish heart regeneration, where macrophages and neutrophils promote epithelial-to-mesenchymal transition (EMT) and epicardial cell expansion ^33,34^. NiCo further identified upregulated genes among these covarying factors (Fa2 and 3 of epicardial-derived cells), which were associated to protein localization to cell periphery, regulation of morphogenesis (Fa2) and cell migration (Fa3) (Fig. S7B). Many Fa3 genes are also known to be involved in EMT and tumor metastasis, such as *Lum, Ecscr, Serpinf1, Sema7a, Fibin,* and *Nes* (Fig. S7C).

TGFβ signaling promotes epicardial cell EMT and motility in mouse and human hearts ^35^. NiCo predicted TGFβ1-TGFβR1 signaling from MP to epicardial cells, where *TgQ1* is highly expressed in Ly6C^high^ and *Spp1^high^* mo/MP, and the receptors *TgQr1, 2* and *3* are all expressed in epicardial-derived cells (Fig. S7D,E). Consistent with this, immunofluorescence staining of Gr-1 (MP and granulocyte marker), mesothelin (MSLN, epicardial-derived cell marker), TGFβ1 and TGFβR2 in day 7 hearts confirmed colocalization of Gr-1+ cells with MSLN+TGFβR2+ cells in the IZ (Fig. 5J). Taken together, these data suggest that early infiltrating Ly6C^high^ and *Spp1^high^* mo/MP may promote EMT of epicardial cells, facilitating cellular plasticity that allows migration into the infarcted myocardium and directly or indirectly promoting epicardial cell transdifferentiation into pericytes for angiogenesis.

### Coordinated regulation of lymphocyte recruitment and activity

On day 7, many lymphoid populations expanded in spatial proximity to CCR7+ DC, cDC2/moDC and lymphatic EC proximal or within the IZ (Fig. 5K,L). We observed an interaction between CCL21a on lymphatic EC and CCR7 on DC ^36^ (Fig. S7F,G). CCR7+ DC expressed *Cxcl16*, which chemoattracts *Cxcr6*+ lymphocytes ^37^, including ILC2, MAIT-like cells, γδT cells and Treg, along with both immune-activation (such as CD86-CD28) and regulatory (such as CD86-CTLA4) signaling (Fig. S7F,G). The IZ-localizing lymphocytes also show spatial proximity to infiltrating myeloid cells such as cDC2/moDC, Ly6C^high^ and *Spp1*^high^ mo/MP, but not to rMP on day 7 (Fig. 5L,M and Fig. S7H). ILC2, MAIT-like cells and effector CD4+ T cells (Th2) were the only cells that expressed *Il4*, an anti-inflammatory cytokine known to induce M2 polarization in MP ^38^ expressing the receptor gene *Il4ra* (Fig. 5N and Fig. S7I). These IL-4+ lymphocytes were proximal to Ly6C^high^, *Spp1^high^* and *Trem2^high^* MP in the day 7 IZ (Fig. 5M).

Infiltrating monocytes can differentiate from a Ly6C^high^ mo/MP state, via a *Spp1^high^Trem2^high^* intermediate state, into *Trem2^high^Gdf15^high^* MP ^20^. Others reported their differentiation into *Cx3cr1^high^*MHCII^high^ rMP instead ^19^. Our pesudotime analysis predicted a sequential progression through Ly6C^high^, *Spp1^high^, Trem2^high^,* to *Cx3cr1^high^/Timd4+* states (Fig. 2G,H and 5O). Since our day 7 spatial data suggested their interaction with IL-4+ lymphocytes (Fig. 5M), we investigated whether IL-4 could drive proinflammatory Ly6C^high^ mo/MP into our sequentially predicted cell states. We cultured bone marrow-derived macrophages (BMDM) ^20^ with cardiac damage-associated molecular patterns (DAMP), i.e. necrotic cells and IFNγ, for 24 hours to mimic the early MI microenvironment (Time point 1). To mimic the day 3-7 MP -IL-4+ lymphocyte interactions, the BMDM were then exposed to IL-4 for 48 hours (Time point 2), followed by cytokine-free medium for 48 hours (Time points 3 and 4). At each time point, BMDM were harvested for FACS of surface markers identified from the pseudotime-gene expression profiles (Methods and Fig. 5P,Q). DAMP-treated BMDM upregulated surface markers of the Ly6C^high^/proinflammatory phenotype (State 1). Subsequent exposure to IL-4 was associated with a reduction in Ly6C^high^ markers and increased proportion of *Spp1^high^*, *Trem2^high^* MP expressing the proliferative marker Mki67 (States 2–3). After IL-4 removal, cells had decreased *Spp1^high^*/*Trem2^high^* MP markers and low Mki67, but upregulated rMP markers (States 4–5) (Fig. 5R and Fig. S7L,M). In summary, our data support a model where transient IL-4 signaling induces a transition of proinflammatory mo/MP into proliferative *Spp1^high^* and *Trem2^high^* states, followed by progression into slow-cycling resident phenotypes.

### Mutual control of proliferation by molecular crosstalk of macrophages and fibroblasts in the lesion

FB and MP are key determinants of cardiac wound healing. On day 7, *Spp1^high^* mo/MP, *Trem2^high^/Cx3cr1*^high^ MP and *Postn+Thbs4+* myoFB are densely enriched in the IZ. The (myo)FB remain colocalized with *Cx3cr1^high^* MP until day 28 but acquired an *Il1rapl1+* transition state. (Fig. 6A,B). Consistent with this, NiCo covariation analysis showed that FB and MP latent factors (Fa3 and Fa2, respectively) covary significantly (Fig. 6C). One highly correlated gene in MP Fa3 is the secretory ligand *Gas6* ^39^, along with complement factors, the mature MP marker *Mrc1*, and growth factor *Igf1*. Genes anti-correlating with this factor are associated with stress response (*Fos*), interferon pathways (*Ifitm3, Isg15*) and proinflammatory marker *Ly6c2* (Fig. 6C). The covarying FB Fa2 is positively correlated with ECM-remodeling genes such as *Mmp2, Lum* and *Dcn*, and anti-correlated with gene sets for cell cycling (Fig. 6D,E) and localization of telomerase RNA to Cajal bodies, (Supplementary Fig. S8A) suggesting a reduction in both cell cycle and telomerase activities. Consistently, our single cell data predicted GAS6-AXL and PROS1-AXL interactions between *Trem2^high^*, *Cx3cr1^high^*/*Timd4+* MP and transition/quiescent FB starting from day 7, when downregulation of growth phase 2-mitotic phase (G2M) cell cycle genes in FB was observed (Fig. 6F-I). Taken together, these data suggest that the interaction between *Trem2 ^high^/Cx3cr1^high^* MP and FB entails silencing of inflammatory roles of the MP, as well as suppression of cell cycle activity in FB.

**Figure 6.**
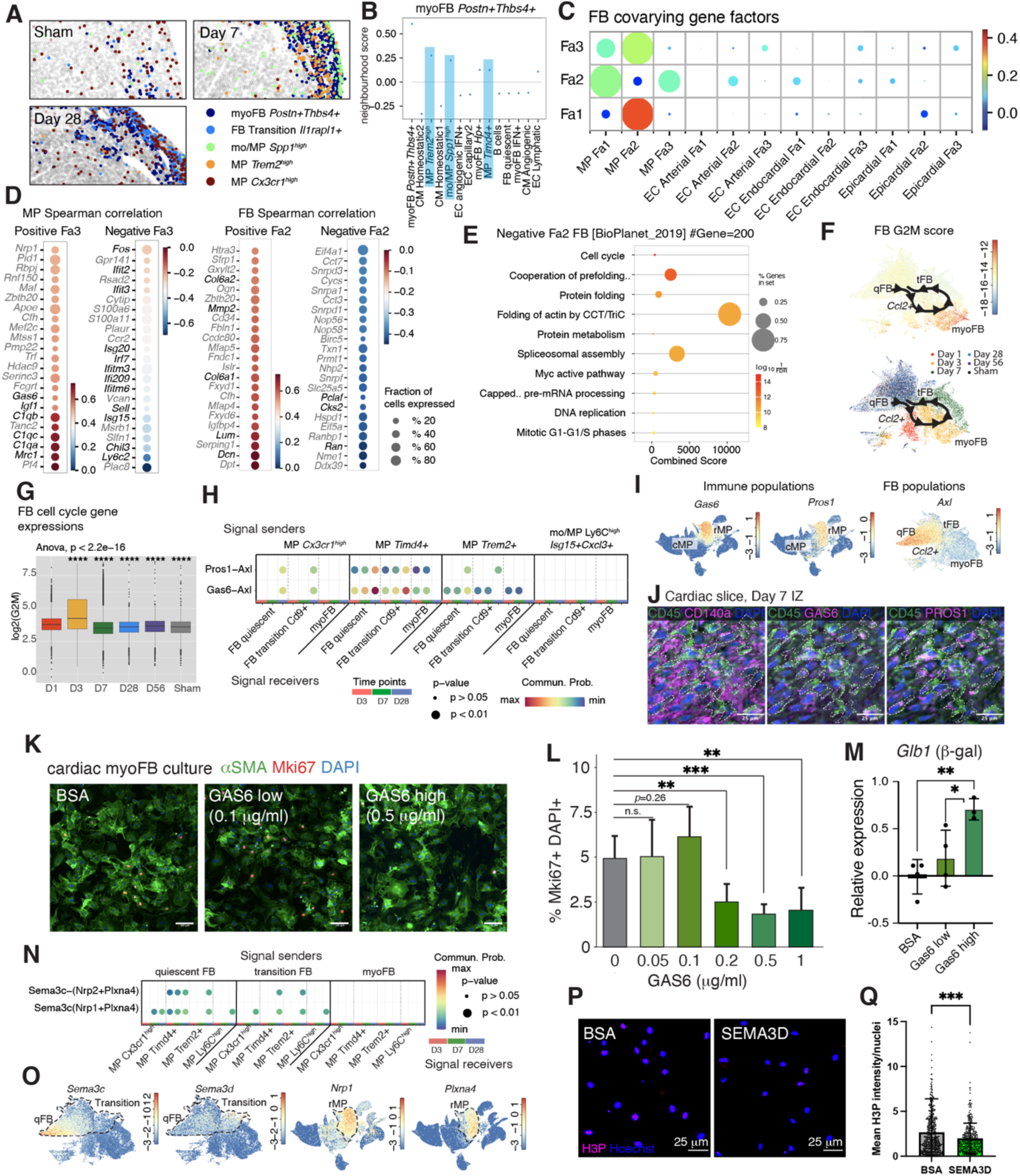
Spatial-neighborhoods of the fibrotic niche. **A**, Spatial maps showing MP and FB subtypes in the wound (LV edge in sham). **B**, day 7 neighborhoods surrounding *Postn+Thbs4+* myoFB, with MP highlighted in blue. **C**, spatial gene factor co-variations of MP neighborhood in day 7 heart. Circle size scales linearly with −log_10_(p-value), and circle color indicates ridge regression coefficients. **D**, genes positively and negatively associated with gene factor 3 (Fa3) of MP and Fa2 of FB, respectively. Highlighted genes in MP-positive-Fa3: secreted ligands; MP-negative-Fa3: early proinflammatory/IFN pathway; FB-positive-Fa2: ECM genes, FB-negative-Fa2: cell cycle. **E**, pathways negatively associated with FB Fa2 (BioPlanet 2019 database). **F**, FB UMAP with inferred cell fate transition and cell cycle G2M gene score (upper), and time points post-lesion. **G**, boxplot comparing log2-transformed aggregated G2M gene expression of FB over time. **H**, CellChat ligand-receptor interaction prediction between MP (signal senders) and FB subtypes (signal receivers) across day 3–28. **I**, immune cell UMAP showing *Gas6* and *Pros1* expression, and FB UMAP showing *Axl* expression. **J**, multiplexed-IF staining of day 7 post-LAD IZ. Dotted lines highlight CD45+ GAS6+ PROS1+ immune cells contacting myoFB (CD140a+). **K – M**, mouse cardiac myoFB culturing experiment (Methods). Cultured myoFB were harvested for IF staining with αSMA and Mki67 (**K**), quantification of % Mki67+ nuclei (DAPI+) in αSMA+ cells (myoFB) (**l**), as well as qRT-PCR quantification of senescence marker gene *Glb1* (β-gal) (**M**). **N**, CellChat prediction for FB (senders) and MP subtypes (receivers), across day 3-28. **O**, UMAPs of FB (left) and immune (right) clusters plotting SEMA3 family and their receptor’s gene expression. **P** and **Q**, mouse BMDM culturing. BMDM were exposed to BSA/SEMA3D for 72 hours, then harvested for H3P IF (**P**) with intensity quantification (**Q**). For (**L**), (**M**), (**P**), (**Q**), statistical significance was calculated by unpaired two-tailed t-test for two experimental groups. * *p* < 0.05, ** *p* < 0.01, *** *p* < 0.001, and **** *p* < 0.0001; ns, not significant.

We validated ligand–receptor co-localization on the protein level by multiplexed-immunofluorescence (IF) on day 7 LAD slices. Consistent with our spatial data, large numbers of FB (CD140a+) and MP (CD45+ GAS6+ PROS1+) were closely packed in the IZ, but not in BZ and RZ (Fig. S8B). IF of TGFβ-activated FB (αSMA+), using FB isolated from 6-week-old mice, showed protein expression of the GAS6/PROS1 receptor AXL (Fig. S8C). To functionally validate this interaction, we performed *in vitro* culture of either non-activated FB or myoFB with recombinant mouse GAS6 ligand and assessed proliferation by Mki67+ nuclei staining (Fig. 6K,L, Fig. S8D-J), which confirmed the proliferation-suppressive effect of GAS6. A similar effect was observed with the PROS1 ligand (Fig. S8H,I). As genes related to telomerase activity were also anti-correlated with FB Fa2, we assessed expression of senescence and cell cycle arrest markers, such as *Glb1* (β-galactosidase), *Trp53* (p53) and *Cdkn1a* (p21). myoFB exposed to high concentration of GAS6 ligands showed activation of most of these genes (Fig. 6M and Fig. S8J), suggesting that GAS6 exposure can trigger cell cycle arrest and cellular senescence.

We next studied ligands expressed by FB that could mediate suppression of proliferation of neighboring *Trem2+/Cx3cr1*^high^ MP, as suggested by anti-correlation of MP Fa3 with activation and proinflammatory phenotypes (Fig. 6C and Fig. 5D). Expression of the Semaphorin family ligand *Sema3d* correlates to FB Fa2 (enriched in tFB and qFB), while its receptors *Nrp* and *Plxna* ^40^ were upregulated in rMPs; this interaction was also predicted by CellChat (Fig. 6N,O and Fig. S8K,L). *In vitro* culture of BMDM with recombinant mouse SEMA3D ligand validated its proliferation-suppressive function, as demonstrated by a lower mean nuclear phospho-histone 3 (H3P) intensity, indicating lower cell cycle activity (Fig. 6P,Q).

Finally, we asked whether this FB-MP co-suppression mechanism is conserved in human hearts. We re-analyzed a recent human MI snRNA-seq dataset ^6^ and focused on the IZ cells (Fig. S8M,N), which comprise a large population of myoFB (*PDGFRA+ ACTA2+ COL1A1^high^*) with high cell cycle gene expression. The myeloid population, which mostly consists of *PTPRC+ ITGAM+* MP, contains both infiltrating *CCL18+* and resident *LYVE1+* phenotypes. Similar to observations in mice, *GAS6* expression is enriched in resident versus circulatory MP, while *AXL* expression is upregulated in FB versus myoFB (Fig. S8O-Q). Exposure of primary human cardiac FB to recombinant human GAS6 (hGAS6) resulted in suppression of proliferation upon addition of hGAS6 in serum-containing medium (Fig. S8R). Similarly, for TGFβ-activated myoFB, proliferation was also suppressed by hGAS6 (Fig. S8S,T). Taken together, our findings confirm suppression of proliferation in activated human FB.

### Synergistic induction of cardiomyocyte dedifferentiation by their niche

CM are collectively responsible for regulating the pumping action of the heart. The regenerative capacity of CM is lost during their postnatal maturation, and CM loss upon injury results in hypertrophy of the remaining CM and fibrosis. Our CM snRNA-seq data (Fig. 7A-D, Fig. S9A,B) encompass large proportions of pre-hypertrophic (*Myh7+ Ankrd1+*) and hypertrophic (*Xirp2+ Nppa+ Nppb+ Ankrd1+ Myh7+*) CM states on days 1 and 3, followed by a transient increase in angiogenic and *Slit2+* states on day 7, expressing angiogenic ligands, growth factors and patterning signals such as *Angpt1, Fgf12, Cntn2, Slit2* and *Slit3*. For the spatial data, the homeostatic, (pre-)hypertrophic, and dedifferentiating states were recovered by spatial DEG, while *Slit2+* and angiogenic states were not distinguished (Fig. S9C,D). Most of these annotated states were distributed across the BZ and RZ (but excluded from IZ/FZ), with enrichment of hypertrophic CM in the BZ (Fig. 7E,F). Hypertrophic CM exhibit gene factor covariations with FB (FB Fa1/2 – hypertrophic CM Fa2) (Fig. S9E,H). Fa2 in hypertrophic CM is positively correlated to genes promoting cardiac muscle contraction, myofibril assembly, and Ca^2+^ release by sarcoplasmic reticulum. *TgQ1*, a known inducer of CM hypertrophy ^41^, is expressed in quiescent FB (Fig. S9G-I). In return, hypertrophic CM express the TGFβ homolog *TgQ2*, which may contribute to FB activation into myoFB as they migrate across the BZ into IZ (Fig. S9G-I).

**Figure 7.**
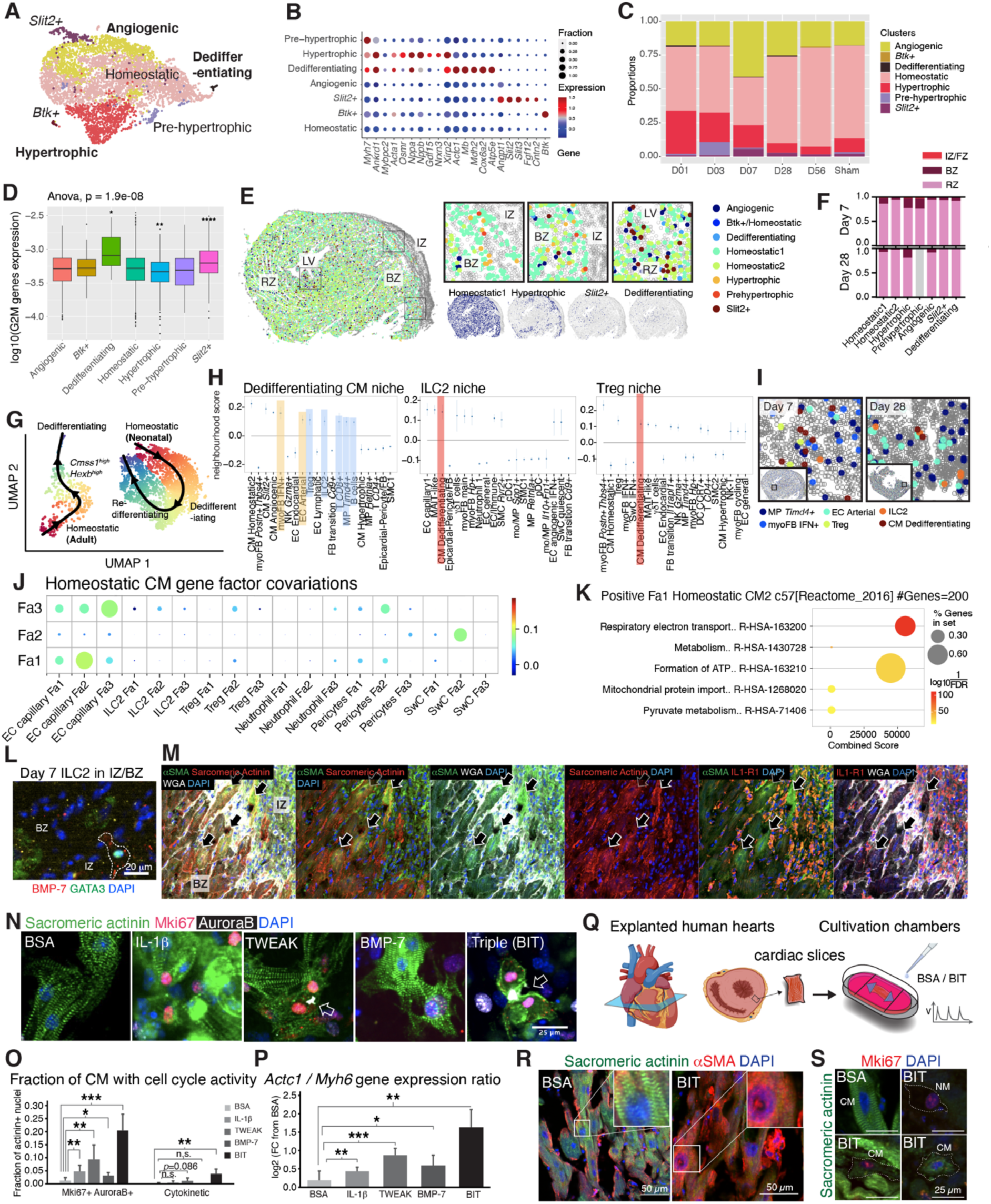
Spatio-temporal interactions of the CM niches. **A**, UMAP of scRNA-seq data for CM subtypes from day 1 to 56 post-lesion. Post-lesion enriched subtypes are highlighted in bold. **B**, DEGs in each subtype. Dot size indicates fraction of cells expressing the gene. **C**, sub type proportion across time. **D**, quantification of summed G2M gene expression on logarithmic scale. **E–F**, annotated spatial maps of CM subtypes on day 7, with detailed annotation scheme (**E**), and quantifications across days 7 and 28 (**F**). **G**, comparison of pseudotime trajectory from homeostatic to the dedifferentiated state in adult versus neonatal CM post-lesion ^4^. **H**, spatial neighborhoods of dedifferentiating CM, ILC2 and Treg on Day 7. **I**, dedifferentiated CM niche visualization on days 7 and 28. **J**, spatial gene factor co-variation of homeostatic CM cluster 2 on day 7. Circle size scales linearly with −log_10_(p-value), and circle color indicates ridge regression coefficients. **K**, pathways positively associated with homeostatic CM cluster 2 Fa1 (GO Reactome 2016 database). **L and M**, IF of day 7 LAD in IZ-BZ for ILC2 (KIT+ GATA3+ BMP-7+) (**L**) and dedifferentiating CM (Sarcomeric actinin+ aSMA+) (**M**). WGA labels cell membranes. Arrows point to IL1-R1+ dedifferentiating CM. **N**, primary P7-CM culturing (see methods). Cells were exposed to ligands for 48 hours. Representative IF images of ligand-exposed CM. CM (Sacromeric actinin+) with Mki67+ AuroraB+ DAPI+ nuclei demonstrate cell cycle activity. White arrows point to cytokinetic morphology indicating CM cell division. **O**, image quantifications of CM with cell cycle activity (left) and undergoing cytokinesis (right) for data in (**N**). **P**, qRT-PCR of *Actc1* (progenitor marker) and *Myh6* (mature marker) of cultured CM. *Actc1*-to-*Myh6* expression ratios (as a progenitor state score) were shown. **Q**, schematic diagram of human cardiac slice culture (see Methods). **R**, representative images showing loss of sacromeric structures in the BIT-exposed CM for human cardiac slice culture. **S**, representative images showing Mki67+ CM and NM in BIT-treated slices, but not in BSA-treated slices. White dotted lines highlight cell boundaries. For (O–P), statistical significance was calculated by unpaired two-tailed t-test for two experimental groups. * *p* < 0.05, ** *p* < 0.01, *** *p* < 0.001, and **** *p* < 0.0001; ns, not significant.

Our data also revealed two rare populations, which were annotated as *Btk+* CM and dedifferentiating CM. The latter shared several markers with hypertrophic CM, e.g., *Myh7* and *Ankrd1*, and were absent in sham. Dedifferentiating CM upregulated progenitor genes (*Actc1*, *Mb* and *Mdh2*), metabolic genes (*Cox6a2* and *Atp5e*) and cell cycle genes, indicating a progenitor-like, proliferation-active state (Fig. 7A-D, Fig. S9J,K). Unlike hypertrophic CM, dedifferentiating CM showed increased glucose metabolism, but decreased cardiac conduction gene expression (Fig. S9M,N), and they were sparsely located in BZ and RZ (Fig. 7E).

Neonatal mice at postnatal day 1 (P1) can regenerate their myocardium after MI, with the emergence of progenitor or dedifferentiating CM subtypes ^4^ represented by a *Mb^high^Myh6^low^* population specific to P1 but absent from P8 hearts post MI (Fig. S9O). By integrating our data with this neonatal CM dataset, we confirmed the resemblance of our adult dedifferentiating CM to neonatal progenitor CM (Fig. S9P,Q).

Pseudotime analysis of neonatal post-MI CM revealed transient activation into a dedifferentiated state, followed by re-differentiation/conversion of these cells back to the homeostatic state (Fig. 7G). In comparison, adult CM could enter the dedifferentiated state via a *Cmss1^high^ Hexb^high^* state. A cocktail of genes encoding different intracellular proteins (such as *Mb, Actc1, Cmss1, Hexb*) and transcription factors (such as *Nfe2l, Jun, Jund, Elf1, Tcf4, Ebf1, Sox7, Cenpa, Hey2*) was sequentially switched on along this trajectory. However, the re-differentiation path was only observed among neonatal CM, but not in adult CM (Fig. 7G, Fig. S9T,U), supporting the notion that adult dedifferentiated CM may not have the full capacity to give rise to mature CM.

Next, we investigated potential niche drivers of CM dedifferentiation. Our day 7 spatial data showed proximity of dedifferentiating CM to ILC2, Treg, CD8+ T cells, B cells, *Timd4+* rMP, IFN+ myoFB, and arterial EC (Fig. 7H,I). Gene expression patterns of EC covaried with homeostatic CM genes, indicating upregulation of respiration and pyruvate metabolism, potentially priming cells towards a dedifferentiating phenotype (Fig. 7J,K). Consistent with our spatial data, IF confirmed the presence of dedifferentiating CM (Sarcomeric actinin+ αSMA+/Myoglobin^high^) and ILC2 (BMP-7+GATA3+) in both the BZ and RZ (Fig. 7L,M and Fig. S9V,W).

To shortlist candidate ligand–receptor interactions, cell surface receptors were extracted from DEG of dedifferentiating CM. After pooling with dedifferentiation trajectory-associated transcription factors (TF) (gene modules 14 and 15 in Fig. S9T), we applied GENIE3 ^42^ to predict a gene regulatory network that connects these receptors and TF that pinpoint key cell fate regulators (Fig. S10A,B). Based on these predictions, TF *Zfp697* may regulate hypertrophic and structural genes (such as *Myh7, Ankrd1* and *Xirp2*), *Nfe2l* and *Jun* may regulate metabolism-related genes (such as *Mdh1, Cox6a2* and *Atp5l*), and *Tcf4* and *Zfand5* may regulate expression of many marker genes that are normally expressed in EC or other vascular cells (such as the receptors *Kdr, Vwf, Pecam1, Cdh5, Kit* and *Pdgfrb*, and the TFs *Sox7, Sox17* and *Sox18*). Some of these interactions involve receptor genes, such as *Il1r1*, *Bmpr2, Fzd3, Tnfrsf12a* and *Ncl.* As dedifferentiating CM exhibit high proximity to ILC2, arterial EC, and *Timd4+* MP, these cell types may provide the corresponding ligands. Although the individual roles of these ligands in promoting CM cell cycle were reported in different animal models, their cellular sources have remained unknown ^8,43,44^. While arterial EC express the TNFRSF12A ligand TNFSF12 (TWEAK), ILC2 express BMP-7 binding to BMPR2, neutrophils express the proinflammatory cytokine IL-1β that binds IL-1R1 (Fig. S10C,D). We confirmed IL-1R1 protein expression in dedifferentiating CM (Fig. 7M).

To functionally assess the role of these ligands in CM cell fate determination, primary CM isolated from P7 mouse LV were exposed to recombinant mouse IL-1β, TWEAK, and BMP-7. As CM are mostly non-regenerative in P7 hearts ^4^, CM proliferation was almost absent when primary CM were exposed to BSA as negative control (<2% actinin+ nuclei were Mki67+ AuroraB+). In comparison, exposure to IL-1β, TWEAK or BMP-7 individually induced a significant (2- to 5-fold) increase in CM cell cycle activity (Fig. 7N,O). Additionally, qRT-PCR revealed a significantly higher level of expression of the CM progenitor marker *Actc1* relative to the mature CM marker *Myh6* (Fig. 7P). Lower expression of the hypertrophic CM marker *Xirp2* was observed when primary cells were exposed to BMP-7 (Fig. S10E), suggesting induction of dedifferentiation and suppression of hypertrophy. Importantly, although the individual effect of each of these ligands on promoting full CM cell cycle was marginal, simultaneous exposure of CM to all three ligands (BIT) resulted in synergistic induction of *Actc1* and proliferation with 10-fold increase in Mki67+ AuroraB+ CM nuclei and significant increase in cytokinetic CM (*p* < 0.01) (Fig. 7O,P). Taken together, these data suggest that the interaction of CM within a niche hosting ILC2, EC, and neutrophils could potentially induce dedifferentiation by co-signaling via the BMP-7–BMPR2, IL1β–IL-1R1, and TWEAK–TNFRSF12a axes.

To further elucidate whether the role of these ligands is conserved in humans, we inspected the presence of dedifferentiating CM in human post-MI snRNA-seq data ^6^. Indeed, a small fraction of CM, found in RZ and IZ, expressed progenitor-like genes, such as *MB, ACTC1, NKX2-5, ATP5PO* and *COX7B*, as in our adult mouse data. These cells are transcriptionally distinct from hypertrophic CM (*XIRP2+*), and are almost absent in the control group, supporting the suggestion that dedifferentiation occurs (Fig. S10F). For the candidate receptors, human CM express low levels of *TNFRSF12a* and *IL1R1*, but higher levels of *BMPR2*, suggesting responsiveness to BMP-7 signaling. Culturing primary human CM progenitor-like cells (isolated from ventricles of failing adult human donor heart) with recombinant hBMP-7 ligand led to strong upregulation of the progenitor marker *ACTC1* (Fig. S10G). To test whether our ligands can induce CM dedifferentiation in native human myocardium, we utilized an *ex vivo* culturing system for cardiac slices excised from explanted hearts of adult human transplant recipients. The slices were cultured in biomimetic chambers and exposed to both preload and regular electrical stimulations. BSA (control) or the candidate ligands (BIT) were added to the culture, and slices were harvested after 6 days (Fig. 7Q). Exposure to BIT ligands led to reduction in actinin structures (Fig. 7R) and increased αSMA expression in CM (Fig. S10H,I), as determined by IF and FACS, respectively. While the control slices contained no Mki67+ proliferative cells, BIT-exposed slices exhibited cell proliferation events both for CM and NM (Fig. 7S and Fig. S7J). In conclusion, CM dedifferentiation niche factors could serve conserved functional roles in both mouse and human myocardium, potentially driving CM cellular plasticity upon cardiac injury.

## Discussion

After MI in the adult heart, replacement of infarcted myocardium with scar tissue achieves a balance between mechanical support to prevent tissue dilatation on the one hand and the inevitable impairment of cardiac physiological function due to the loss of electro-mechanically functional myocardium on the other.

Achieving this balance involves complex cellular communication networks that coordinate the proper responses of multiple cell types across tissue locations and stages of remodeling. Currently available cardiac single-cell sequencing and spatial datasets provide insights into cell type-specific responses ^6,45–47^, but typically lack coverage of the dynamics of interacting cell types in the lesion. Moreover, most murine MI studies have collected the entire LV for sequencing, diluting the representation of relevant cell types from the lesion area in the dataset, thereby complicating interpretation of the resulting data. Available spatial transcriptomic data were generated with low-resolution sequencing-based techniques. Although deconvolution can be applied to infer the cell type composition of each pixel or spot ^48^, more subtle cell state modulation and covariation of gene expression between cell types within the same spot cannot be inferred. Thus, there is a need for a comprehensive post-lesion cardiac atlas that maps spatio-temporal dynamics of cell type architecture in the wound and thereby supports identification of the key heterocellular interactions that shape the outcome of cardiac remodeling.

By densely sampling all stages of post-lesion remodeling, harvesting tissue from the lesion area, and enriching rare cell populations, we were able to overcome previous limitations and create an atlas with unprecedented resolution of the cell types involved in cardiac tissue repair. We not only captured the behavior of cell types known to play a critical role in scar formation (such as FB and MP), but we also uncovered the dynamics of poorly characterized rare cell types, such as unconventional T cells, innate lymphocytes, SwC, epicardial-derived cells, and dedifferentiating CM. These discoveries open the door to more in-depth studies that will improve our understanding of the role of these cell types after lesioning or in homeostatic hearts. The similar damage response we observed for Cryo and LAD suggests that both models are suitable for investigating the role of these cell types in cardiac healing.

The integration of sequencing data with single-cell resolution spatial transcriptomics through our novel NiCo algorithm ^18^ enabled the identification of novel signaling interactions in lesion niches and their downstream effects on cell states.

Our spatio-temporal analysis identified an immune-fibrotic niche in the IZ, composed of multiple mo/MP subtypes, myoFB and lymphocytes. Th2, ILC2 and MAIT-like cells produce IL-4 on day 3-7 which promotes M1 (Ly6C^high^) MP transdifferentiation into *Spp1^high^/Trem2^high^* states. At later stages, when lymphocyte numbers are reduced, decreased IL-4 signaling promotes further transdifferentiation of *Trem2^high^* MP into resident-like phenotypes. Additionally, both the intermediate (*Trem2^high^*) and some resident (*Cx3cr1^high^*) MP interact with myoFB in the niche, promoting mutual suppression of proliferation of both cell types in the maturing scar through GAS6/PROS1 and SEMA3 signaling (Fig. S10K).

GAS6 and PROS1 are vitamin K-dependent proteins that bind TAM receptors, including TYRO3, AXL, and MER ^49^. GAS6 is generally known as a mitogenic ligand ^39,50^, whereas an opposing role has been reported for PROS1 ^51^. However, in the case of cardiac FB, we observed both GAS6- and PROS1-mediated cell cycle reduction, which is associated with cellular senescence. It remains an open question which additional factors determine whether cell cycle activity is positively or negatively regulated in response to vitamin K-dependent protein signaling.

Our data further revealed niche interactions that support CM dedifferentiation (Fig. S10L), a poorly understood process with great potential to promote functional recovery of the heart. Unlike hypertrophic CM, which are highly enriched in the BZ and exchange TGFβ signaling with migrating FB, dedifferentiating CM were sparse throughout the myocardium, consistent with previous studies ^5,7,52^. Our data indicate that the simultaneous activation of multiple signaling pathways (BMP-7, TWEAK, and IL-1β) is required to synergistically push cells into the dedifferentiated state. ILC2 are the main source of BMP-7 in the dedifferentiation niche but are very rare in the adult myocardium. γδT cells and Treg also express BMP-7 but occur at low frequency in adult myocardium, and they do not show significant spatial interactions with CM. These findings suggest the importance of lymphocytes, especially unconventional populations, for potential cardiac regeneration.

Importantly, understanding how to promote effective CM dedifferentiation may confer substantial therapeutic potential for the restoration of adult cardiac function post MI in the future. Through our experimental data on isolated human primary CM progenitors and *ex vivo* cultured human cardiac slices, we have demonstrated that our identified niche ligands may promote human CM dedifferentiation.

We also identified a stage-dependent neighborhood of epicardial-derived cells exhibiting elevated EMT gene signature when interacting with Ly6C^high^ and *Spp1^high^* mo/MP via TGFβ signalling (Fig. S10M), potentially differentiating into pericytes and FB. Indeed, the differentiation potential of these cells is still under debate, and they may contribute to the CM fate ^53^. The spatio-temporal dynamics of epicardial-derived cells is therefore an exciting topic for future studies which may lead to new interventions for promoting *de novo* CM generation. In conclusion, this spatial-temporal cell type atlas for cardiac injury is a valuable resource that will serve as a reference for understanding relevant heterocellular interactions during cardiac repair.

## Methods

### Data and code availability

Single-cell RNA-seq data have been deposited at GEO. Accession number is publicly available as of the date of publication.

Source code used in this study is available on Github at: https://github.com/Andyson-Chan/myocardial_infarction/tree/main

### Animal experiments

All animal experiments were carried out according to the guidelines in Directive 2010/63/EU of the European Parliament on the protection of animals used for scientific purposes; they were approved by the local authorities in Baden–Württemberg, Germany (Regierungspräsidium Freiburg, G21-129), and by the animal welfare officer of the Centre for Experimental Models and Transgenic Services-Freiburg (X21-03R; X23-03R).

### Mouse cardiac surgery

LV cryoablation (Cryo), ischemia-reperfusion injury after left anterior descending artery (LAD) ligation, and sham surgery were performed on female wildtype C57BL/6J mice at 12 weeks of age. Surgery and Cryo were performed as previously described ^54^. In brief, mice were anaesthetized by intraperitoneal injection of 80–100µL of anesthesia solution (20mg/mL ketamine (Ketaset; Zoetis, Parsippany-Troy Hills, NJ, USA), 1.4mg/mL xylazine hydrochloride (0.12% Rompun; Bayer, Leverkusen, Germany), in 114mM NaCl (0.67% m/v; B. Braun Melsungen, Melsungen, Germany)). After induction of deep anesthesia, eye ointment (Bepanthen containing 50mg/mL dexpanthenol; Bayer) was applied, 500µL glucose solution (278mM glucose = 5% (m/v); B. Braun Melsungen) was injected intraperitoneally, and 250µL of analgesia solution (10µg/mL buprenorphine (Temgesic; Indivior Inc., North Chesterfield, VA, USA) in 154mM NaCl (0.9% m/v, B. Braun Melsungen)) was injected subcutaneously in the neck. Mice were shaved on the left side of the thorax (precordial region) and on the right leg, placed on a warming platform of a small animal physiology monitoring system (Harvard Apparatus, Holliston, MA, USA), and the front extremities were fixed with tape. A rectal thermometer was inserted to allow regulation of the heating platform to ensure a body temperature of 37°C. Thermometer and tail were fixed with tape. Mice were intubated for positive pressure ventilation (Kent Scientific, Torrington, CT, USA; 40% O_2_, 120 breathing cycles per minute). Isoflurane was supplied at 5% until the animal stopped spontaneous respiratory movements, and then reduced to 2.0–2.5%. An infrared blood oximeter was attached to the right leg to track hemoglobin oxygen saturation, ventilation was adjusted if needed. The surgical field was disinfected using Softasept N (B. Braun Melsungen). Skin and muscles were cut along the third intercostal space, and a rib spreader was used to separate the ribs. The pericardium was cut, and the epicardial surface was dry-blotted using a cellulose pad.

For mice undergoing ventricular cryoablation, a hexagonal metal probe (stainless steel, 2.5mm edge-normal distance) was pre-chilled in liquid nitrogen and applied for 8–10s to the free LV mid-wall, positioned to avoid major coronary vessels. After retraction of the probe, the time until the tissue regained a deep-red color was recorded (consistently within 5–10 seconds). For mice in which the ischemia-reperfusion injury was performed, the LAD was identified and ligated with an 8-0 suture, while a narrow plastic tubing was inserted below the suture. Successful ligation was indicated by the infarcted area changing in color, from red to grey. After 30–45 minutes, the tubing and suture were removed. For sham surgery, application of the metal probe and ligation was not performed. The rib spreader was removed, and the thorax closed using a 6-0 silk suture around the third and fourth ribs (4–5 single knots). Before final closure, any remaining air was removed from the thorax using a small cannula. Isoflurane application was stopped, and the skin was closed with a 4-0 silk suture. Once the mouse started breathing spontaneously, intubation and fixation were terminated, and the mouse was transferred to a heated and oxygenated wake-up chamber. Analgesia was maintained for 72 hours post surgery via twice-daily subcutaneous injection of 250µl of buprenorphine (10µg/mL in 154mM NaCl, injected in the morning and late afternoon). During the night, buprenorphine was also supplied via the drinking water (10µg/mL buprenorphine (Subutex lingual tablets, Indivior) in 20mM glucose solution).

### Tissue Processing and Cell Isolation

The tissue collection and slicing protocol was adapted from a previously published protocol ^54,55^. In brief, mice received sodium-heparin solution (16units/g body weight) by intraperitoneal injection, before being sacrificed by cervical dislocation five minutes later. Their chests were then opened and their hearts were excised. To wash out the blood from the heart, hearts were cannulated and flushed with cold ‘cutting solution’ (in mM): NaCl 138, NaH_2_PO_4_ 0.33, KCl 5.4, MgCl_2_ 2, HEPES 10, glucose 10, CaCl_2_ 0.5, 2,3-butanedione 2-monoxime (BDM) 30, pH-adjusted to 7.3 with 1 M NaOH at 37°C, osmolality 330 ± 10mOsm/L. Tissue blocks of LV free wall containing the post-Cryo or post-LAD lesion centre (identified as a region appearing mostly white), BZ and adjacent myocardium (or corresponding tissue areas from sham animals) were excised and embedded in low-melting-point agarose (4% (m/v)) at 37°C, and then put on ice. Agarose blocks were glued to the stage of a precision vibratome and cut into 300µm thick slices (60Hz cutting frequency, 1.5Hz amplitude, model 7000smz-2; Campden Instruments, Loughborough, UK). The resulting tissue slices were subjected to chemical fixation, cryopreservation, or cell isolation.

For chemical fixation, slices were exposed for 30 minutes to 4 % paraformaldehyde (PFA)-containing phosphate-buffered saline (PBS) at room temperature (RT), washed three times in PBS, and then stored in PBS at 4°C.

To cryopreserve tissue, each slice was placed flat in the bottom center of a cryo-mold (Tissue-Tek, 4566), immersed in a thin layer of optimal cutting temperature compound (OCT, Tissue-Tek, 4583). To fix the orientation and keep the slice flattened, an additional cryomold of the same size was placed on top of the tissue, before snap-freezing in liquid nitrogen. Frozen slices were stored at −80°C until cryosectioning.

For cell isolation, tissue slices were stored at 4°C in cutting solution and warmed to RT before initiating cell isolation. Tissue slices were transferred to ‘enzymatic solution’ containing (in mM): KCl 20, KH_2_PO_4_ 10, MgCl_2_ 2, Taurin 20, glucose 10, L-glutamic acid potassium salt monohydrate 100, BDM 30, osmolality 310 ± 10, pH adjusted to 7.3 with KOH at 37°C. Tissue was digested for 12 minutes with proteinase XXIV (concentration: 0.5mg/mL; Sigma-Aldrich, St. Louis, MO, USA) and further digested with Liberase TL Research Grade (concentration: 0.23mg/mL with 5μM CaCl_2_; Hoffmann-La Roche, Basel, Switzerland) for up to 45 minutes. To remove large tissue fragments, the cell suspension was filtered through a nylon mesh (pore size: 1mm × 1mm).

### NM antibody labelling and FACS sorting

FACS sorting was performed using a BD FACS Symphony S6 or FACSAria III Cell Sorter (BD BioSciences, Franklin Lakes, NJ, USA). The same gating and collection strategy was applied across all time points post lesioning to ensure comparable cell type ratios. For each of the collected NM sample for 10x GEM generation, isolated cells were first blocked by the TruStain FcX Fc blocking antibody (Biolegend, 101319) for 15 minutes at 4°C. Cells were labeled for 30 minutes at 4°C in the dark with fluorophore-conjugated antibodies against various surface markers, including CD45-Pacific Blue (Biolegend, 103125), Ter119-APC/Cy7 (Biolegend, 116223), CD31-PE (Biolegend, 102507), CD146-PE (Biolegend, 134703), CD14-FITC (Biolegend, 123307) and CD11c-APC (Biolegend, 117309), together with the Zombie-NIR viability dye (BioLegend, 423105), which labels dead cells. All antibodies were diluted at 1:100 in FACS buffer [PBS with 10% fetal bovine serum (FBS, VWR, MDTC35-016-CV) containing 1% Penicillin-Streptomycin solution (P/S, 100U/mL, Gibco, 15140122)]. When performing FACS gating, half of the cells were collected from the Zombie-NIR and Ter119-negative gate (unbiased), and the other half were collected from the enrichment gates (See Supplementary Figure 1a). Cells were sorted into FACS buffer-coated 1.5mL tubes, each with 40μL of FACS buffer with 1:1000 diluted Murine RNase Inhibitor (New England Biolabs, M0314L). After collection, cell counting was performed. In each sample, 10,000 cells from the unbiased tube and 10,000 cells from the enrichment tube were pooled together for 10x GEM generation.

### Single nucleus isolation from mouse hearts

For nucleus isolation, vibratome-cut cardiac slices were placed in MACS M-tubes (Miltenyi Biotec, 130-093-236), containing 1mL of nuclear staining buffer [PBS with 1% BSA (Miltenyi Biotec, 130-091-376) and 1:500 diluted (0.2U/μL) Murine RNase Inhibitor, pre-cooled to 4°C. The tube was placed into a gentleMACS Octo dissociator (Miltenyi Biotec, 130-096-427), and then dissociated using the included “Protein_01.01 M-tube” protocol. The M-tubes then underwent pulse-spinning, were flushed with 1mL of nuclear staining buffer, and were filtered through a 40μm cell strainer (Falcon, 1172689). The eluates were collected in 50mL Falcon tubes, which were then centrifuged for 5 minutes at 500*g*, 4°C, brake 5. Supernatants were removed, and pellets were resuspended in 500μL of nuclear staining buffer.

### Nuclei antibody labelling and FACS sorting

Isolated nuclei from different time points were first labelled with 1:100 diluted αPCM-1 antibody (MERCK, HPA023370) in nuclear staining buffer for 5 minutes at 4°C (not on ice), then with 1:100 diluted fluorophore-conjugated anti-Rabbit IgG secondary antibody (Biolegend, 406419) for an additional 25 minutes at 4°C in the dark. Samples then underwent centrifugation (5 minutes, 500*g*, 4°C, brake 5), followed by multiplexing with CellPlex reagents according to the manufacturer’s instruction (10x Genomics, PN-1000261), resulting in time point-specific labeling in each sample. After labeling, nuclei were washed with 2ml nuclear staining buffer and centrifuged (5 minutes, 500*g*, 4°C, brake 5). Labelled nuclei from all time points were then pooled together in 1mL nuclear staining buffer containing 10μg/mL Hoechst 33,342 (Thermo Fisher Scientific, H3570), and underwent FACS sorting to enrich for Hoechst+ PCM-1+ nuclei. Enriched nuclei were collected in a 1.5mL tube pre-coated with nuclear staining buffer, containing 150μL of nuclear staining buffer with 1:100 diluted of Murine RNase Inhibitor. Within the collected nuclei, 10μL was stained with Trypan Blue solution (ThermoFisher, 15250061) at a ratio of 1:1, and was visually inspected under a light microscope (20x objective) to assess nuclei shape and morphology. If the majority of nuclei appeared to be intact (i.e., with no visible disruption of nuclear membrane), this was followed by nuclear counting. 20,000 nuclei were used for 10x GEM generation.

### Sample processing for sequencing

In each sample, up to 20,000 sorted cells were loaded into GEM Chip (10x Genomics, PN-1000127) and encapsulated into emulsion droplets using the Chromium Controller (10x Genomics, Pleasanton, CA, USA) according to the manufacturer’s instruction. cDNA library generation and amplification were performed using the “Chromium Next GEM Single Cell 3ʹ Reagent Kits v3.1 (Dual Index)” protocol according to manufacturer’s instructions (10x Genomics, PN-1000268 and PN-1000190). The prepared cDNA library was then sequenced on the Nextseq2000 platform (Illumina, San Diego, CA, USA). The CellRanger v6.1.1. pipeline was used to generate a digital gene expression matrix starting from raw data. For alignment and quantification of gene expression the reference transcriptome has been built using the mouse mm10-2020-A as reference genome. De-multiplexing of CellPlex barcodes into individual samples was performed by Seurat ^48^.

### scRNA-seq and snRNA-seq data analysis

Single-cell clustering and data analysis were performed using VarID2.^56^. Each sample condition was initially represented by one raw gene count matrix. Cells within each matrix were labelled with their condition. The labelled matrices were merged and processed into a single cell object via the “SCseq” function. Only cells with more than 1,000 UMI counts were considered for clustering and analysis. Mitochondrial genes, ribosomal genes, and predicted genes with Gm-identifier were filtered (FGenes argument in filterdata function). Analysis of k-nearest neighbor was performed using the “pruneknn” function. For the clustering combining all cells across all conditions, batch effect correction was performed using the harmony package ^57^. In summary, the following settings were used “large=TRUE,regNB=TRUE,knn=25,no_cores=8,seed=12345, FSelect = T,batch = batches, bmethod = “harmony”. To define clusters, “graphCluster” function was applied, with Leiden clustering ^58^. Settings were as follows: “pvalue=0.01, use.weights = T, use.leiden = T, leiden.resolution = 1”. Gene expression values were normalized by correcting variability associated with total transcript count per cell by a negative binomial regression ^56^, which could be visualized in heatmaps of gene expression UMAPs. For Boxplot gene expression representation, UMI counts were normalized by dividing transcript counts in each cell by the total transcript count per cell before multiplying by the minimum total transcript count across all cells. For cell type annotations, two schemes were applied throughout this study. The first scheme defined all identified cell types from the main clusters (of all cell types) in detail, and annotated all subtypes and states within each major cell types (such as all lymphocyte populations and all FB cell states), provided that each annotated cell (sub)type had distinct marker gene expressions. Known cell (sub)types were annotated according to published names, and newly identified types were annotated based on their enriched marker gene(s). This is referred to as the “detailed annotation scheme”, and yielded 78 cell types in total. Alternatively, a “simplified annotation scheme” was also defined, where some cellular subtypes without distinctive separations in UMAP were pooled together to form one major type, resulting in only 38 cell types. The simplified annotation scheme was only used for the spatial neighborhood interaction analysis. To analysis specific cell types in a higher resolution, cell identities from clusters of CM, FB, vascular cells, immune cells, and SwC were collected. The raw UMI counts from these cells were extracted from the count matrices and underwent re-clustering, with the same parameters described above. Within the immune cell subclusters, cells from all lymphocyte clusters (except B cells) were collected, and further re-clustered into a lymphocyte-only subcluster.

### Differential expression analysis

Differential expression analysis was computed with the “diffexpnb” function of the RaceID3 (v0.2.5) algorithm. Detection of differentially expressed genes between specific groups of cells was performed with a method similar to that previously reported ^59^. In brief, a negative binomial distribution, which captures the gene expression variability for each group of cells, was inferred based on a background model of the expected transcript count variability estimated by RaceID3 ^60^. Based on the inferred distributions, a P value for the significance of the transcript counts between the two groups of cells was estimated and multiple testing was corrected for using the Benjamini–Hochberg method.

### Pathway enrichment analysis

Pathway enrichment analysis was performed with the “enrichPathway” function from the ReactomePA R package ^61^, with a P value cutoff of 0.05. Inputs were ENTREZ gene IDs of genes selected by differential expression analysis or otherwise specified. Gene concept networks were constructed using the enriched pathways from the two cell populations under comparison. Pathways were located at the center node of each cluster, while the corresponding genes were shown as connected smaller nodes. The size of each pathway node corresponded to the number of contributing genes.

### Lineage trajectory inference

Transition probabilities between two clusters were calculated as the geometric mean of the individual link probabilities connecting the two clusters. To order cells pseudo-temporally, the Slingshot method ^29^ was applied on a desired dimensionally reduced RaceID object (selected cell clusters). Using the FateID package, pseudo-time expression profiles were derived by self-organizing maps (SOM) and grouped into modules.

### Gene regulatory network inference for CM

To infer gene regulatory networks, the R Bioconductor package GENIE3 ^42^ was applied with default parameters on the full dataset. For the inference of CM dedifferentiation gene regulatory network, genes from SOM-identified module 14 and 15 were used as the input gene list, excluding genes expressed in the mitochondria. Candidate regulators were defined as transcription factors in the gene list, plus differentially expressed cell surface receptor genes (dedifferentiating CM against other CM, FC > 1). Only the first 120 links were reported in the plotted map.

### Ligand-receptor interaction analysis

To predict potential ligand-receptor interactions using the single-cell and nuclear datasets, the CellChat package ^62^ was applied. Briefly, this tool first identified enriched ligand and receptor genes across each manually annotated cell types (*p* < 0.05). A detailed annotation scheme (as defined above) was used for interaction analysis. Using CellChat’s own ligand-receptor database, predicted interactions between defined cell types, conditions (time points), and pathways was visualized via the “netVisual_bubble” function. This dot plot shows communication probability (indicated by the color of each dot) and the corresponding *p*-value (the size of each dot) for each predicted interaction.

### Reanalysis of published human MI snRNA-seq datasets

Seurat objects from 3 post-MI patient IZ samples were downloaded from the Zenodo data archive (https://zenodo.org/records/6578047), and annotated as “IZ_P21_ext1”, “IZ_P22_ext2”, and “IZ_P23_ext3”. After clusters of myeloid cells and FB were confirmed by marker genes, corresponding cells were re-clustered by VarID2 ^56^ with the same parameters as above. For FB subclustering UMAP construction (compumap), the parameters used were spread = 6, min_dist = 0.5. For myeloid cell subclustering UMAP construction, the parameters used were spread = 1, min_dist = 0.5.

### snRNA-seq data comparison with published neonatal mouse MI datasets

The neonatal MI CM atlas ^4^ was downloaded from NCBI Gene Expression Omnibus, with accession number GSE130699. Briefly, the count matrices from all time points (P1 mice MI 1 and 3 days; P1 mice sham: 1 and 3 days; P8 mice MI 1 and 3 days; P8 mice sham: 1 and 3 days) were collected, with cells from each condition being labelled. The resulting matrices, together with our adult CM post-cryoablation count matrices, underwent clustering by VarID2 ^56^, with the same settings and parameters described above. For the UMAP construction (compumap), the parameters used were min_dist = 3.5, spread = 4.

### Spatial transcriptomics gene panel design

The Xenium Mouse Tissue Atlassing Panel (379 genes) was applied and topped up with 96 custom genes based on DEGs of each cell (sub)type identified in our scRNA-seq data. The probes had two complementary sequences to bind the target RNA and contained a third region with a gene-specific barcode. This allowed the ends to ligate into a circular DNA probe for in situ amplification, ensuring high specificity by preventing ligation during off-target binding.

### Spatial transcriptomics sample processing, imaging and preprocessing

All experimental steps followed the guidelines of the Xenium workflow ^63^. Briefly, 5μm formalin-fixed paraffin-embedded tissue sections (transverse plane of the LV) were mounted on a Xenium slide within the desired imaging region. The mRNA was exposed by deparaffinization and permeabilization. A gene panel, as described above, was applied to the sample in 10nM, and targeted mRNA molecules of the 475 target genes. In addition to the gene panel, two negative controls were added to assess non-specific binding and ensure the signals were from binding to RNA and not genomic DNA. Hybridization was performed at 50°C overnight, followed by a washing step to remove unhybridized probes. Additionally, barcode oligo-conjugated antibodies (ATP1A1, E-Cadherin, CD45, αSMA, Vimentin and 18S rRNA marker) labeling the cell boundaries and interior proteins were added to label cellular structures. Samples were then ligated at 37°C for 2 hours for annealing of rolling circle amplification (RCA) primer, followed by circularized probe amplification at 4°C for 1 hour, and 37°C for 2 hours, resulting in amplification of the gene-specific barcode for each RNA binding probe, enhancing the signal-to-noise ratio. Background fluorescence was chemically quenched.

Sections were placed in an imaging cassette and loaded onto the Xenium Analyzer instrument combining a high numerical aperture and a fast area scan camera with a low read noise sensor (achieving a lateral resolution of 200nm per pixel). Images were acquired in sequential hybridization-imaging cycles. In each cycle, unique fluorescently labeled secondary probes targeting RCA-amplified probe-RNA complexes were hybridized, and visualized in combination with 4ʹ,6-diamidino-2-phenylindole (DAPI) staining. Stained samples were imaged, followed by stripping off the secondary probe and starting another round of hybridization and imaging. Z-stacks were acquired at a 0.75μm step size across the tissue thickness.

Images were pre-processed with the built-in pipeline, where low quality signals were filtered out, and the identities of transcripts were decoded using a Xenium codebook. For quality scoring, a Phred-style Q-Score was assigned to each decoded transcript, reflecting confidence in transcript identity. Q-Scores were calibrated using negative control codewords, probes targeting non-biological sequences, and unassigned codewords. Only transcripts with a Q-Score ≥20 were included in downstream analyses.

Multimodal cell segmentation was also performed using the Xenium’s built-in pipeline, in which cells were segmented based on nuclear (DAPI), cell boundary, and interior RNA signals.

### Spatial transcriptomics data analysis

We utilized the NiCo algorithm ^18^ to perform integrative analysis of spatial transcriptomics and scRNA-seq/snRNA-seq data.

*Spatial cell type annotation:* We used all the default parameters for the cell type annotation task, except for the spatial guiding Leiden clustering resolution, which was set to 0.8 for Day 7 and Day 28 samples, and to 0.3 for sham.

*Spatial neighborhood composition analysis:* Analysis was conducted for each cell type with the NiCo interaction module, determining colocalization scores for all niche cell types using logistic regression classifier. A simplified cell type annotation scheme was applied for plotting. Threshold values of 0.14 for day 7, 0.18 for day 28, and 0.20 for sham were used to plot cell type niche interaction maps.

*Niche covariation analysis:* Analysis was conducted for each cell type independently with the covariation module of NiCo using integrated NMF to identify cell type-specific latent factors. NiCo can be used to conduct ridge regression between the latent factors of the central cell type as dependent variables and latent factors of co-localized neighborhood cell types as independent variables. Significant regression coefficients reflect statistical dependence, or covariation, of latent factors belonging to co-localized cell types. We used the default parameters for the identification of ligand-receptor pairs associated with covarying cell type-specific latent factors, and performed pathway enrichment analysis for all genes correlating to each latent factor to infer the associated gene programs.

### Mouse primary CM culturing

Neonatal mouse hearts were harvested from P7 mice (at least 6 hearts were harvested for a single experiment) and were transferred to a 6cm Petri dish (VWR, 734-0006, 353003) containing PBS. Remaining blood was pumped out of the hearts by compression, using forceps to apply gentle pressure. Remaining non-ventricular tissue was removed and the ventricles were minced into small fragments using dissecting scissors. For tissue digestion, a Neonatal Heart Dissociation Kit (Miltenyi Biotec, 130-098-373) was used. Briefly, PBS was replaced by2.5mL of enzyme mix (prepared according to the manufacturer’s instruction). The petri dish was then incubated at 37°C with gentle agitation (65rpm, Incu-Shaker Mini, Z763578-1EA) for 45 minutes. To enhance dissociation rate, the tissue-enzyme mix was pipetted 20 times through a wide-bore 1000µL pipette tube every 15 minutes. After completion of the digestion process, 5mL FACS buffer was added and the mixture was then passed through a 70µm cell strainer (Falcon, 352350). Filtered cells were collected in a 50mL Falcon tube, which was centrifuged at 350*g* for 5 minutes at 4°C. The supernatant was removed, and the pellet was resuspended with 2mL of ACK lysis buffer (Biozym, 882090-FFM) for 5 minutes at 4°C, followed by the addition of 3mL of FACS buffer and re-centrifugation. To magnetically label NM (not CM), the pellet was resuspended with 100µL of 1:10 FACS buffer-diluted neonatal cardiomyocyte isolation antibody cocktail (Miltenyi, 130-100-825) for 15 minutes at 4°C in the dark. For magnetic selection, the mixture was topped up with 500µL FACS buffer, and then loaded into an MS column (Miltenyi Biotec, 130-042-201) in a MACS magnetic separator (Miltenyi Biotec, 130-042-102), pre-loaded with FACS buffer. The column was washed with 3 × 500µL FACS buffer. All the flow-through was collected and combined. Collected cells (CM) were counted. 10,000 CM per well were seeded onto a 96-well plate (ThermoFisher, 164588) precoated with 0.1% Galectin (InSCREENex, INS-SU-1015-50mL) in 200µL full IMDM medium (composed of IMDM (Gibco, 12440053) supplemented with 10% FBS, 1X MEM Non-Essential Amino Acids Solution (NEAA, Gibco, 11140050) and 1% P/S solution). After plating, CM were incubated at 37°C in a 5% CO_2_ cell culture incubator overnight. The next day, medium was replaced by full IMDM medium supplemented with 100ng/mL of either Recombinant Mouse IL-1β (Biolegend, 575102), Recombinant Human TWEAK (Biolegend, 566402), Recombinant Mouse BMP-7 Protein (R&D Systems, 5666-BP-010/CF), or BSA. Cells were incubated at 37°C, 5% CO_2_ for 48 hours, and were harvested for downstream quantification.

### Mouse primary cardiac FB and myoFB culturing

Ventricles were harvested from 6-week-old mice. Ventricular tissue was minced into small fragments using dissecting scissors. Heart extraction and cell isolation followed the adult heart dissociation protocol described above. To enrich cardiac FB, MojoSort was performed. Briefly, isolated cells were centrifuged (300*g*, 5 minutes, 4°C), resuspended in 1:100 FACS buffer-diluted Biotin-conjugated CD140a antibody (ThermoFisher, 13-1401-82), and incubated for 15 minutes at 4°C. Cells were then centrifuged (300*g*, 5 minutes, 4°C), after which the supernatant was discarded and 100µL of MojoSort^TM^ Streptavidin Nanobeads (1:10 diluted in FACS buffer, Biolegend, 480016) was added to the mixture; this was then incubated for another 15 minutes at 4°C in the dark. After incubation, the mixture was transferred to a 5mL FACS tube, and inserted into a MojoSort Magnet (Biolegend, 480019), pre-cooled with ice, for 5 minutes. The supernatant was discarded. The tube was removed from the magnet, and the cells remaining in the tube were resuspended with 0.5mL FACS buffer. After cell counting, the cell solution was diluted by adding full DMEM medium (DMEM (DMEM, high glucose, GlutaMAX™ Supplement, pyruvate, Gibco, 31966021), 10% FBS and 1% P/S solution) and seeded into a 96-well plate, with 10,000 cells per well in 200µL medium. After plating, cells were incubated at 37°C in a 5% CO_2_ cell culture incubator overnight. The next day, the medium was removed and cells were washed twice with PBS, before being incubated with serum-free DMEM with Recombinant Mouse GAS6 protein (0.05– 1000ng/mL, R&D Systems, 986-GS-025/CF), Recombinant Mouse Protein S/PROS1 (100ng/mL, R&D Systems, 9740-PS-050/CF), or an equivalent amount of BSA. Cells were incubated at 37°C in 5% CO_2_ for 48 hours, and were then harvested for downstream quantification. For myoFB culturing experiments, plated FB were first activated for differentiation into myoFB using 100ng/mL TGFβ overnight in medium with 10% FBS, followed by recombinant mouse GAS6 or BSA exposure in serum-free and TGFβ-free medium for 48 hours. Cells were then harvested for downstream quantification, including IF qPCR and FACS analysis.

### Human primary cardiac FB culturing

Human cardiac fibroblasts (HCF) were purchased from Promocell (C-12375), having been isolated from the ventricles of adult human hearts and cryopreserved. Cells were defrosted, resuspended in full HCF medium (Growth Medium 3 with supplements, Promocell, C-23025), and allowed to grow in a T75 flask (Merck, C7231-120EA) until 80% confluence. HCF were then trypsinized using 3mL of Trypsin-EDTA (0,25%) with phenol red (Gibco, 25200072) for 5 minutes at 37°C, followed by centrifugation (300*g*, 5 minutes, 4°C) and pellet resuspension by HCF medium. After cell counting, cells were plated in a 96-well plate, with 5,000 cells per well in 200µL medium. Thereafter, cells were incubated at 37°C in a 5% CO_2_ cell culture incubator overnight. For human cardiac myoFB generation, cells were exposed to 0.1µg/mL of Recombinant Human TGF-β1 (Biolegend, 781802). The next day, the medium was removed. Cells were washed twice with PBS, and then incubated with either serum-free DMEM or full DMEM medium, with the addition of Recombinant Human Gas6 protein (R&D Systems, 885-GSB-050) or the equivalent amount of BSA. Cells were incubated at 37°C, 5% CO_2_ for 48 hours, and were harvested for downstream quantification.

### Human primary progenitor-like CM culturing

Human progenitor-like CM isolated from the ventricles of adult human hearts were purchased from Promocell (C-12810). Cells were defrosted, resuspended in full myocyte growth medium with supplements (Promocell, C-22070) and allowed to grow in 37°C, 5% CO_2_ until 80% confluency. After medium removal, CM were then trypsinized using 3mL of Trypsin-EDTA (0.25%) with phenol red (Gibco, 25200072) for 5 minutes at 37°C, followed by centrifugation (300*g*, 5 minutes, 4°C) and pellet resuspension in myocyte growth medium. After cell counting, cells were seeded into a 96-well plate, with 10,000 cells per well in 200µL medium. After plating, cells were incubated at 37°C in a 5% CO_2_ cell culture incubator overnight. The next day, cells were washed with fresh medium with 0.1µg/mL recombinant human BMP-7 protein (Biolegend, 595601) or an equivalent amount of BSA. Cells were incubated at 37°C, 5% CO_2_ for 48 hours, and then harvested for downstream quantification.

### Generation of danger associated molecular patterns (DAMP) from murine heart

For the generation of DAMP, murine hearts from C57BL/6 mice were collected and pulverized in a mortar under liquid nitrogen. The powder was then resuspended in PBS supplemented with protease inhibitor cocktail at a concentration recommended by the manufacturer (Sigma, 4693159001), using three hearts per mL. DAMP were stored at −20°C until further use.

### BMDM culture and polarization

Bone marrow was isolated as described previously ^64^. Briefly, bone marrow cells were isolated as follows. The femur and tibia bones of C57BL/6 mice were flushed with cold PBS. Cells were centrifuged, and the pellets were resuspended in single-cell suspension in MP medium containing DMEM (Gibco, 41965-039) supplemented with 10% fetal calf serum (Sigma, 12133), 100units/mL penicillin (Gibco, 15140122), 100units/mL streptomycin (Gibco, 15140122), 1mM L-glutamine (Gibco, 25030081), 1x non-essential amino acids (Gibco, 11140-035), 1x sodium pyruvate (Gibco, 11360-070) and 10-15% L929 supernatant (SN). L929 SN was generated as previously described ^65^, with the necessary concentration determined via bioassays. Non-adherent progenitors of MP were harvested after 24 hours of cultivation and seeded at a concentration of 0.5 x 10^6^ per mL for six days in MP medium. 50% of medium was renewed every other day. For polarization, cells were treated with 1% DAMP and 50ng/mL IFNγ in MP medium for 24 hours. After 24 hours, the medium was removed and exchanged with MP medium containing 10ng/mL IL-4. After 48 hours, the medium was removed again, and cells were treated with un-supplemented MP medium for 24 hours or 48 hours. Throughout culturing, cells were kept at 37°C in 5% CO_2_.

### Flow cytometry analysis with Cytek

Flow cytometry analysis was performed as previously described ^66^. Briefly, the cells were incubated on ice with Zombie NIR (BioLegend, 423105) for 10 minutes in PBS. After washing with PBS containing 0.5% BSA, cells were stained with fluorophore-conjugated antibodies (see Table) for surface antigens for 35 minutes on ice in PBS containing 0.5% BSA. Cells were then fixed with Foxp3/TF staining buffer following the manufacturer’s recommendations (eBioscience, 00-5523-00) for 35 minutes on ice. For intracellular staining, cells were incubated with fluorophore-conjugated antibodies (see Table) targeting intracellular antigens for 35 minutes at RT in 1x permeabilization buffer (eBioscience, 00-8333-56). Sample data acquisition was performed at the Aurora Flow Cytometer (Cytek), and analyzed with Tree Star (FlowJo LLC) and Prism (GraphPad Software Inc).

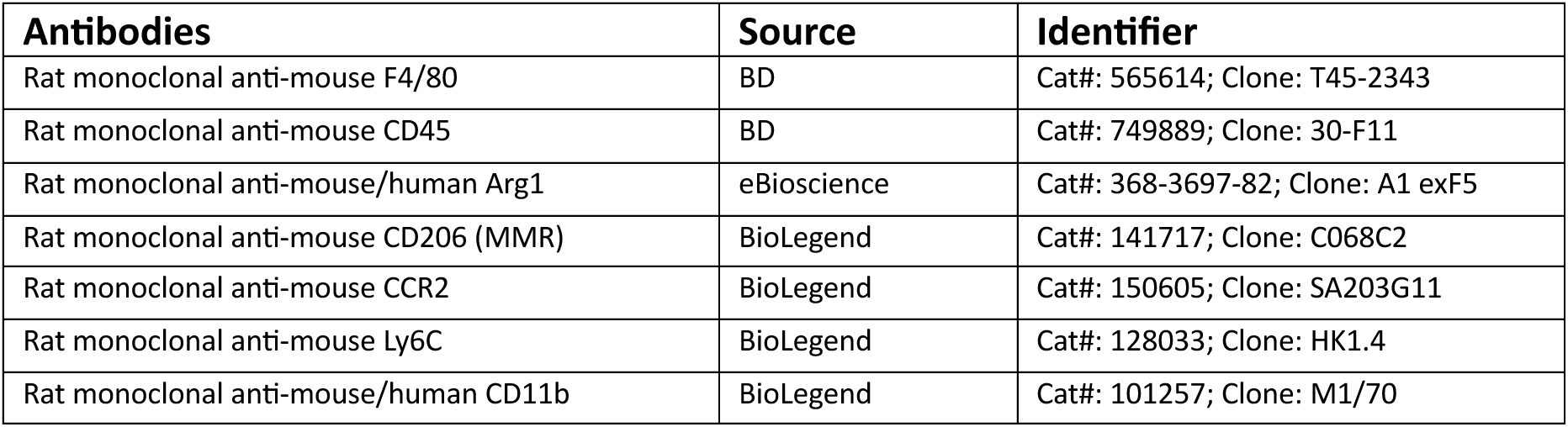
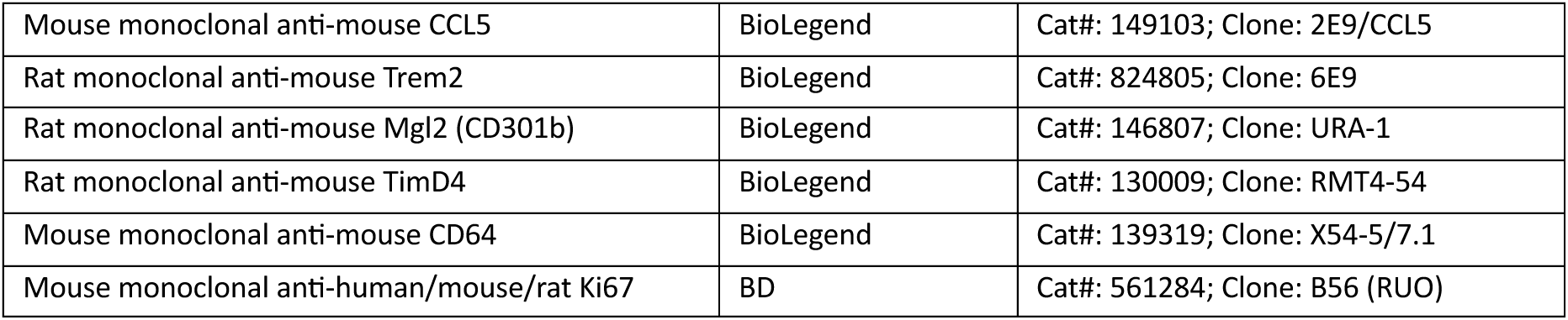

| **Antibodies** | **Source** | **Identifier** |
| --- | --- | --- |
| Rat monoclonal anti-mouse F4/80 | BD | Cat#: 565614; Clone: T45-2343 |
| Rat monoclonal anti-mouse CD45 | BD | Cat#: 749889; Clone: 30-F11 |
| Rat monoclonal anti-mouse/human Arg1 | eBioscience | Cat#: 368-3697-82; Clone: A1 exF5 |
| Rat monoclonal anti-mouse CD206 (MMR) | BioLegend | Cat#: 141717; Clone: C068C2 |
| Rat monoclonal anti-mouse CCR2 | BioLegend | Cat#: 150605; Clone: SA203G11 |
| Rat monoclonal anti-mouse Ly6C | BioLegend | Cat#: 128033; Clone: HK1.4 |
| Rat monoclonal anti-mouse/human CD11b | BioLegend | Cat#: 101257; Clone: M1/70 |
| Mouse monoclonal anti-mouse CCL5 | BioLegend | Cat#: 149103; Clone: 2E9/CCL5 |
| Rat monoclonal anti-mouse Trem2 | BioLegend | Cat#: 824805; Clone: 6E9 |
| Rat monoclonal anti-mouse Mgl2 (CD301b) | BioLegend | Cat#: 146807; Clone: URA-1 |
| Rat monoclonal anti-mouse TimD4 | BioLegend | Cat#: 130009; Clone: RMT4-54 |
| Mouse monoclonal anti-mouse CD64 | BioLegend | Cat#: 139319; Clone: X54-5/7.1 |
| Mouse monoclonal anti-human/mouse/rat Ki67 | BD | Cat#: 561284; Clone: B56 (RUO) |

### SEMA3D assay

Upon differentiation, murine BMDM were treated for 72 hours with 100ng/mL SEMA3D (R&D Systems, 9386-S3-025/CF) in PBS and PBS containing 0.5% BSA as a control, respectively. Cells were fixed in 4% PFA, washed, and treated with 1 x ReadyProbes Mouse-on-Mouse IgG Blocking Solution (Invitrogen, R37621) according to the manufacturer’s instruction. Mouse anti-histone H3 antibody (Abcam, ab14955, 1:100) was then incubated overnight at 4°C. Cells were washed and goat anti-mouse IgG Alexa Fluor 647 secondary antibody (Invitrogen, A32728, 1:1000) was incubated for 1 hour at RT. Nuclei staining was performed with Hoechst 33342 according to the manufacturer’s instructions. Fluorescence intensity measurements were performed on a Leica TCS SP8 X inverted confocal microscope (SCI-MED imaging facility).

### Human cardiac slices culture

Transmural LV human cardiac samples (N=2) were obtained from failing hearts during LV assist device implantation (LVAD), according to protocols approved by the institutional ethics committees of the University of Erlangen-Nürnberg and the University of Leipzig. Studies followed the guidelines of the extended Declaration of Helsinki. All patients (or their legal guardians) gave their written informed consent.

Preparation and culture of LV slices was carried out as previously described ^67,68^. M199 (Sigma, M4530) was used as a culture medium and supplemented with β-mercaptoethanol (50μM), 1:100 penicillin/streptomycin (Biochrom, A2213), cortisol (50nmol/L), insulin (10ng/mL), transferrin (5.5μg/mL) and selenite (6.7ng/mL). Cardiac slices were mounted into specialized culture chambers (MyoDish, MD-1.2, InVitroSys) and diastolic force was set to 1.5mN. Slices were continuously paced at 0.5Hz by electrical field stimulation. Medium was partially exchanged every 48 hours. After 16 days of culture, slices were randomly divided into two groups. The control group (n=5) was treated with 1% FBS. The BIT group (n=6) was treated with a combination of BMP-7, human recombinant TWEAK, and human recombinant IL-1β at a concentration of 0.1µg/mL each. Accounting for 0.2mL of evaporated volume, 1.6mL of medium was removed every 48 hours and 1.8mL of fresh medium containing the three substances or FBS was added. After six days of treatment, the slices were fixed in 4% PFA solution for 45 minutes and then stored in PBS at 4°C.

### Multiplexing immunohistochemistry (IHC) and immunofluorescence (IF) staining

For IF staining of mouse cultured CM or cardiac FB cultures, cells in 96-well plates were first washed with PBS, then fixed with 4% PFA in PBS (Sigma-Aldrich, 252549) for 15 minutes at RT, followed by 2 5-minute washes with 0.3% TritonX-100 in PBS. Cells were then exposed to 70% ethanol at −20°C for 1-2 hours. Cells were then returned to RT, washed with PBS (twice, for 5 minutes each time), and blocked in blocking solution (3% BSA and 0.0025% TritonX-100 in PBS) for 1 hour, followed by antibody labelling [1:200 diluted rat Mki67-FITC antibody (Biolegend, 151211). For FB staining, 1:100 diluted rat CD140a-PE (ThermoFisher, 12-1401-81), and 1:250 diluted mouse αSMA primary antibody (Invitrogen, 14-9760-80) were added. For CM staining, 1:200 diluted human Actinin(Sarcomeric)-PE (Miltenyi, 130-123-996) and 1:400 diluted mouse Aurora B primary antibody (BD Bioscience, 611082) were added at 4°C in the dark for 16–24 hours. Cells were washed twice with PBS for 10 minutes, and then incubated with 1:400 goat anti-mouse IgG-AF647 (Biolegend, 405322) and DAPI solution for 2 hours at RT in the dark, followed by two 10-minute washes with PBS. For HCF staining, Mki67-FITC and αSMA primary antibody were used at the same concentrations. For IHC staining of post-lesion cardiac slices, mounted cryosections underwent similar fixation, wash, and antibody incubation steps as for IF. The following antibodies were used: mouse PDGFRa (R&D Systems, AF1062-SP), rat CD45 (Biolegend, 103108), goat GAS6 (R&D Systems, AF986-SP), rabbit PROS1 (Invitrogen, PA5-106880), goat AXL (R&D Systems, AF854-SP), mouse BMP-7 (NovusBio, NBP2-52425), rabbit GATA3 (Cell Signaling Technology, 5852T), goat KIT (R&D Systems, AF1356-SP). For multiplexing, tissues were first imaged after labeling with the first round of antibodies, followed by two rounds of antibody stripping by glycine-HCl buffer [0.1 M Glycine hydrochloride (Merck, G2879-100G), 0.1% Triton X-100, pH 2.2] for 15 minutes. Tissues were then twice washed with PBS for 5 minutes, and then were re-incubated with another round of antibodies.

### Spinning disk confocal imaging and analysis

IXplore SpinSR Olympus super resolution imaging system (Evident, Tokyo, Japan) was used for imaging of IF and IHC samples, with a 20× objective. Image acquisition was performed with the cellSens software (Evident, Tokyo, Japan). 405, 488, 561 and 640nm lasers were used to excite DAPI, FITC, PE and AF647 fluorophores, respectively. The acquisition time was 200ms for each color channel. After acquisition, images were analyzed using Fiji ImageJ, including adjustments to brightness and contrast, followed by cell density quantifications.

### Flow cytometry analysis with FACSCelesta for isolated cardiac FB

Isolated cells well placed in 1.5mL tubes and centrifuged (350*g*, 5 minutes, 4°C) to remove supernatants. Pellets were resuspended in 4% PFA in PBS, and were fixed for 10 minutes at RT, followed by centrifugation (800*g*, 5 minutes, 4°C) to remove the fixative. Cells were washed with PBS containing 0.1% TritonX-100, recentrifuged and then resuspended in DAPI-containing solution (1μg/mL) for 10 minutes in the dark, recentrifuged and resuspended with PBS in FACS tubes (Falcon, 38025). Readout of DNA content was performed by BD FACSCelesta Cell Analyzer (BD BioSciences, Franklin Lakes, NJ, USA), where histograms of DAPI signal intensities across all cells were recorded. Quantitative analysis of the DNA content was performed with the FlowJo v10 software (FlowJo LLC).

### RNA extraction, cDNA synthesis and quantitative PCR (qPCR)

RNA extraction and purification were performed with the PicoPure RNA Isolation Kit (Applied Biosystems, KIT0204) according to the manufacturer’s instructions. cDNA synthesis was performed with RevertAid H Minus First Strand cDNA Synthesis Kit (Thermo Scientific, K1632), with the following components: 4μL RNA, 1μL Random Hexamer, 7μL H_2_O, 4μL 5x Reaction Buffer, 1μL RiboLock RNase Inhibitor, 2μL dNTP mix, and 1μL RevertAid H Minus Reverse Transcriptase. The reaction was performed in thermocycler under the following conditions: 25°C for 5 minutes; 42°C for 60 minutes; 70°C for 5 minutes; 4°C on hold. qPCR of the cDNA samples were performed as follows: 2μL 1:4 diluted cDNA, 5μL iTaq™ Universal SYBR Green Supermix (BioRad, 1725120), 0.6μL 10μM forward and reverse primers, and 2.4μL nuclease-free H_2_O. For mouse and human cardiac FB samples, *Gapdh* was used as a reference gene. For human cardiac FB samples, *GAPDH* was used as a reference gene. For mouse CM samples, *Smchd1* was used as a reference gene as *Gapdh* exhibited variation in expression levels across different subtypes in the snRNA-seq data, while expression of *Smchd1* was more consistent across all CM subtypes. The qPCR reaction was performed with CFX Connect Real-Time PCR Detection System (BioRad, 1855201) for 45 amplification cycles.

### Quantification and Statistical Analysis

All experiments were performed using randomly assigned mice. Statistical significance was calculated by unpaired two-tailed t-tests for two experimental groups, or one-way ANOVA for multiple comparisons. Statistical analyses and quantifications were performed in Excel (Microsoft Corporation), Prism 10 (GraphPad Software Inc), FlowJo (FlowJo LLC), and Fiji imageJ. Figures display means ± SD as indicated. *p* < 0.05 is considered statistically significant. In all figures, * indicates *p* < 0.05, ** indicates *p* < 0.01, *** indicates *p* < 0.001, and **** indicates *p* < 0.0001 (ns indicates “not significant”).

### Additional resources

The sc/snRNA-sequencing and spatial transcriptomics datasets are available for exploration and visualization at: https://www.wuesi.medizin.uni-wuerzburg.de/cardiac_spatiotemporal_atlas/

## Acknowledgments

The authors thank Pia Iaconianni, Stefanie Perez-Feliz, and Simone Nübling (IEKM Freiburg) for performing mouse surgeries and supporting experiments; Konrad Schuldes and Sebastian Hobitz (Flow Cytometry and DNA Sequencing Facility in Max Planck Institute of Immunology and Epigenetics) and Christian Linden (Institute of Systems Immunology) for supporting FACS sorting and analysis; Josef Madl (Microscope Facility Sci-MED, Universitäts-Herzzentrum Freiburg) for imaging support of BMDM experiment; Julieta Aprea Perez (Dresden-concept Genome Center) for performing the Xenium runs; the group of Christoph Kuppe for providing expert advice on spatial transcriptomics; James O’Reilly for writing advices; Reyna Rosales Alvarez and Alexander Dallmann for programming advices; and Judith Schaf, Nadine Vornberger and Serli Kopar for their experimental and technical support. This work was supported by the German Research Foundation (DFG) SFB1425 (#422681845) to D.G., E.R.Z., P.K. and F.S.W., and INST 93/1072-1 (#471222118) to D.G., SPP1937 GA 2129/2-2 to D.G., and two Emmy-Noether fellowships (#396913060, #412853334) to F.S.W. and E.R.Z., by the ERC (818846 — ImmuNiche — ERC-2018-COG) to D.G., and by the Bundesministerium für Bildung und Forschung (BMBF) (TissueNet - 031L0311A to D.G. and CureFib - 01EJ2201C to D.G.). Further funding was provided by the DFG SFB 1525 #453989101 to M.V.

## Author contributions

D.G. conceived, designed, and supervised the study. A.S.F.C. designed, optimized, and performed cell sorting and scRNA-seq. F.S.W and P.K. planned and supervised the mouse heart surgery experiments, and tissue harvest. E.R.Z., J.G., and T.B. optimized and performed cardiac slice sectioning and cell isolation for scRNA-seq. L.M., A.S.F.C. and W.L.C. coordinated and performed the tissue sectioning for spatial transcriptomics. A.S.F.C. and D.G. analyzed and interpreted the scRNA-seq data. A.S.F.C., A.A. and D.G. analyzed and interpreted the spatial transcriptomic data. A.S.F.C. and K.S. designed BMDM culturing experiments, K.S. and A.K. performed the experiments, and M.V. supervised the experiment. A.S.F.C. designed and performed FB and CM validation experiments. A.S.F.C. and W.L.C. performed multiplexed immunofluorescence imaging. A.S.F.C. and L.M. performed data analysis. T.S. and Z.I. performed human cardiac slice culturing of FB validation experiments. A.S.F.C performed staining, imaging and analysis. H.H. constructed the Shinyapp web interface. A.S.F.C and D.G. wrote the manuscript.

## Declaration of interests

DG serves on the scientific advisory board of Gordian Biotechnology.

## Abbreviation

BIT: BMP-7 – IL-1β – TWEAK cocktail
BMDM: Bone Marrow-Derived Macrophages
BMP: Bone Morphogenetic Protein
BSA: Bovine Serum Albumin
BZ: Border Zone
cMP: Circulatory Macrophages
CM: Cardiomyocytes
Cryo: Cryoablation
CTLA-4: Cytotoxic T-Lymphocyte Associated Protein 4
DAPI: 4’,6-Diamidino-2-Phenylindole
DEG: Differentially Expressed Genes
EC: Endothelial Cells
ECM: Extracellular Matrix
EMT: Epithelial-Mesenchymal Transition
FB: Fibroblasts
FBS: Fetal Bovine Serum
FZ: Fibrotic Zone
G2M: Growth phase 2/Mitosis Phase
GAPDH: Glyceraldehyde 3-Phosphate Dehydrogenase
GAS6: Growth Arrest-Specific 6
HIF-1α: Hypoxia-Inducible Factor 1-Alpha
IL-1β: Interleukin 1 Beta
IL-4: Interleukin 4
IL-5: Interleukin 5
ILC2: Innate Lymphoid Cell Type 2
IHC: Immunohistochemistry
IFNγ: Interferon Gamma
IZ: Ischemic Zone
LAD: Ligation of the Left Anterior Descending Artery
LV: Left Ventricle
MAIT: Mucosal-Associated Invariant T cells
mg: Milligram
mL: Milliliter
mo/MP: Monocytes/Macrophages
MP: Macrophages
myoFB: Myofibroblasts
ng: Nanogram
NM: Non-Myocytes
OCT: Optimal Cutting Temperature
OXPHOS: Oxidative Phosphorylation
PDGFRA: Platelet-Derived Growth Factor Receptor Alpha
PROS1: Protein S
qFB: Quiescent Fibroblasts
qRT-PCR: Real-Time Quantitative Reverse Transcription - Polymerase Chain Reaction
rMP: Resident Macrophages
RZ: Remote Zone
scRNA-seq: Single-cell RNA Sequencing
SMC: Smooth Muscle Cells
SwC: Schwann Cells
TCR: T cell Receptor
tFB: Transition Fibroblasts
TGFβ: Transforming Growth Factor Beta
Treg: Regulatory T Cells
tSNE: t-distributed stochastic neighbor embedding
TWEAK: TNF-related Weak Inducer of Apoptosis
µL: Microliter
µg: Microgram
UMAP: Uniform Manifold Approximation and Projection
XIRP2: Xin Actin Binding Repeat Containing 2

