## Supplementary Figures for "Spatio-temporal dynamics of the fibrotic niche in cardiac repair"

Chan et al.

Supplementary Figures 1-10

#### Figure S1

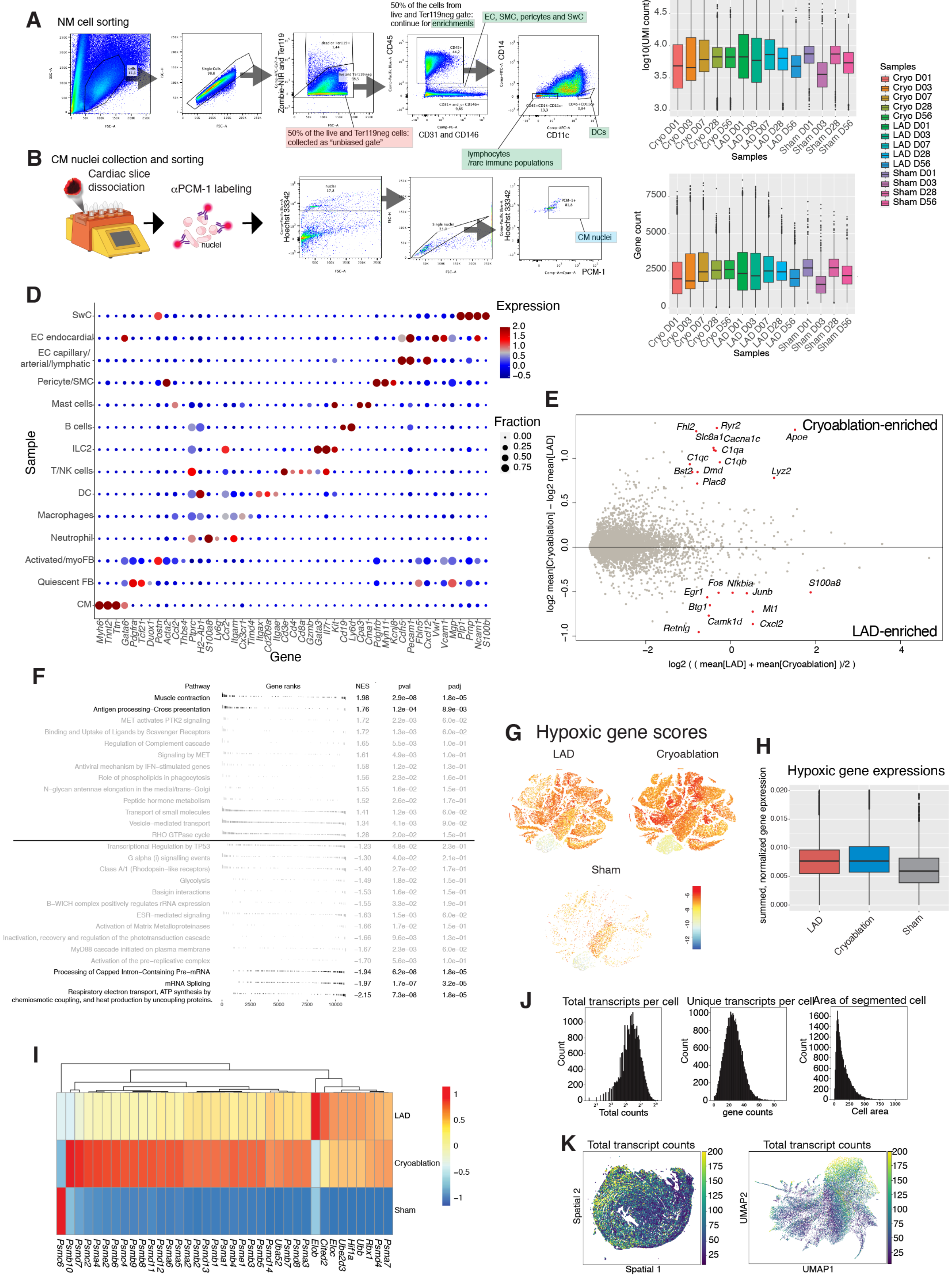

A
NM cell sorting
Zombie-NIR and Ter119
CD45
CD14
50% of the cells from live and Ter119neg gate: continue for enrichments
EC, SMC, pericytes and SwC
50% of the live and Ter119neg cells: collected as "unbiased gate"
CD31 and CD146
CD11c
DCs
lymphocytes /rare immune populations
B
CM nuclei collection and sorting
Cardiac slice dissociation
αPCM-1 labeling
nuclei
Hoechst 33342
CM nuclei
PCM-1
C
log10(UMI count)
Gene count
Samples
Cryo D01
Cryo D03
Cryo D07
Cryo D28
Cryo D56
LAD D01
LAD D03
LAD D07
LAD D28
LAD D56
Sham D01
Sham D03
Sham D28
Sham D56
D
Expression
Fraction
Sample
SwC
EC endocardial
EC capillary/ arterial/lymphatic
Pericyte/SMC
Mast cells
B cells
ILC2
T/NK cells
DC
Macrophages
Neutrophil
Activated/myoFB
Quiescent FB
CM
Myh6 Tnnt2 Ttn Gata6 Pdgfra Tcf21 Duox1 Postn Acta2 Ccl2 Thbs4 Ptprc H2–Ab1 S100a8 Ly6g Ccr2 Itgam Cx3cr1 Timd4 Itgax Cd209a Itgae Cd3e Cd4 Cd8a Gzmb Gata3 Il7r Kit Cd19 Ly6d Cpa3 Cma1 Pdgfrb Myh11 Kcnj8 Cdh5 Pecam1 Fbln5 Cxcl12 Vwf Vcam1 Mgp Plp1 Prnp Ncam1 S100b
Gene
E
Cryoablation-enriched
LAD-enriched
Fhl2 Ryr2 Slc8a1 Cacna1c Apoe C1qa C1qc C1qb Bst2 Dmd Lyz2 Plac8
Fos Nfkbia S100a8 Egr1 Junb Btg1 Mt1 Camk1d Cxcl2 Retnlg
log2 mean[Cryoablation] – log2 mean[LAD]
log2 ( ( mean[LAD] + mean[Cryoablation] )/2 )
F
Pathway | Gene ranks | NES | pval | padj
Muscle contraction | 1.98 | 2.9e–08 | 1.8e–05
Antigen processing–Cross presentation | 1.76 | 1.2e–04 | 8.9e–03
MET activates PTK2 signaling | 1.72 | 2.2e–03 | 6.0e–02
Binding and Uptake of Ligands by Scavenger Receptors | 1.72 | 1.3e–03 | 6.0e–02
Regulation of Complement cascade | 1.65 | 5.5e–03 | 1.0e–01
Signaling by MET | 1.61 | 4.9e–03 | 1.0e–01
Antiviral mechanism by IFN–stimulated genes | 1.58 | 1.2e–02 | 1.3e–01
Role of phospholipids in phagocytosis | 1.56 | 2.3e–02 | 1.6e–01
N–glycan antennae elongation in the medial/trans–Golgi | 1.55 | 1.6e–02 | 1.5e–01
Peptide hormone metabolism | 1.52 | 2.6e–02 | 1.7e–01
Transport of small molecules | 1.41 | 1.2e–03 | 6.0e–02
Vesicle–mediated transport | 1.34 | 4.1e–03 | 9.0e–02
RHO GTPase cycle | 1.28 | 2.0e–02 | 1.5e–01
Transcriptional Regulation by TP53 | –1.23 | 4.8e–02 | 2.3e–01
G alpha (i) signalling events | –1.30 | 4.0e–02 | 2.1e–01
Class A/1 (Rhodopsin–like receptors) | –1.40 | 2.7e–02 | 1.7e–01
Glycolysis | –1.49 | 1.8e–02 | 1.5e–01
Basigin interactions | –1.53 | 1.6e–02 | 1.5e–01
B–WICH complex positively regulates rRNA expression | –1.55 | 3.3e–02 | 1.9e–01
ESR–mediated signaling | –1.63 | 1.5e–03 | 6.0e–02
Activation of Matrix Metalloproteinases | –1.66 | 1.7e–02 | 1.5e–01
Inactivation, recovery and regulation of the phototransduction cascade | –1.66 | 9.6e–03 | 1.3e–01
MyD88 cascade initiated on plasma membrane | –1.67 | 2.3e–03 | 6.0e–02
Activation of the pre–replicative complex | –1.70 | 5.6e–03 | 1.0e–01
Processing of Capped Intron–Containing Pre–mRNA | –1.94 | 6.2e–08 | 1.8e–05
mRNA Splicing | –1.97 | 1.7e–07 | 3.2e–05
Respiratory electron transport, ATP synthesis by chemiosmotic coupling, and heat production by uncoupling proteins. | –2.15 | 7.3e–08 | 1.8e–05
G
Hypoxic gene scores
LAD
Cryoablation
Sham
H
Hypoxic gene expressions
summed, normalized gene expression
LAD
Cryoablation
Sham
I
LAD
Cryoablation
Sham
Psmc6 Psmb10 Psmb7 Psma2 Psme2 Psmb6 Psmc4 Psmd8 Psmd11 Psmd12 Psma6 Psma5 Psmd13 Psmb1 Psma1 Psmb4 Psmd3 Psmb5 Psmd14 Ube2i Uba52 Psmb7 Psmd9 Psma3 Cited2 Elob Eloc Ube2d3 Hif1a Ubb Rbx1 Psmd4 Psma7
J
Total transcripts per cell
Unique transcripts per cell
Area of segmented cell
Count
Total counts
gene counts
Cell area
K
Total transcript counts
Spatial 2
Spatial 1
UMAP2
UMAP1

**Figure S1. Cell type enrichment, gene expression quantifications and comparison of lesion models**

**A**, FACS enrichment and sorting of NM cells. The same gating and collection strategy has been applied across all time points post-lesion to ensure comparable cell type ratios. For each of the collected NM samples for 10x GEM generation, half of the cells were collected from the Zombie-NIR and Ter119-negative gate (highlighted in red), and the other half of the cells were collected from the enrichment gates (highlighted in green). **B**, Preparation and FACS enrichment of CM nuclei. Cardiac slices were dissociated by gentleMACS dissociator with the protocol "Protein_01.01 M tube", followed by washing and DAPI and $\alpha$PCM-1 antibody labeling. Labelled nuclei then underwent FACS sorting to enrich for DAPI+ PCM-1+ nuclei, followed by 10x GEM generation. **C**, boxplots showing overview of transcript counts (UMI) and gene counts from each collected sample (CM + NM). Cryo, cryoablation. **D**, dot plot showing expression z-scores of marker genes that represent different major cell types. Dot diameter represents the fraction of cells of that cell type expressing the plotted gene. **E and F**, comparison of gene expression ($p < 0.05$, highlighted in red) (**G**) and pathways (**H**) that were differentially enriched in cryoablation and LAD samples. Pathway comparison was performed using fast gene set enrichment analysis (fgesa, Korotkevich et al, 2016). Only pathways written in dark were significantly enriched (adjusted $p < 0.05$). **I – K**, comparison of hypoxia-related genes across LAD and Cryo samples. **I**, UMAPs showing summed expression of hypoxic genes in LAD, Cryo and Sham cells and nuclei. **J**, quantified normalized sum of hypoxic gene expression across the 3 conditions. **K**, heatmap comparing the enrichment of each hypoxic gene across the 3 condition ($-1 < \text{score} < 1$).

### Figure S2

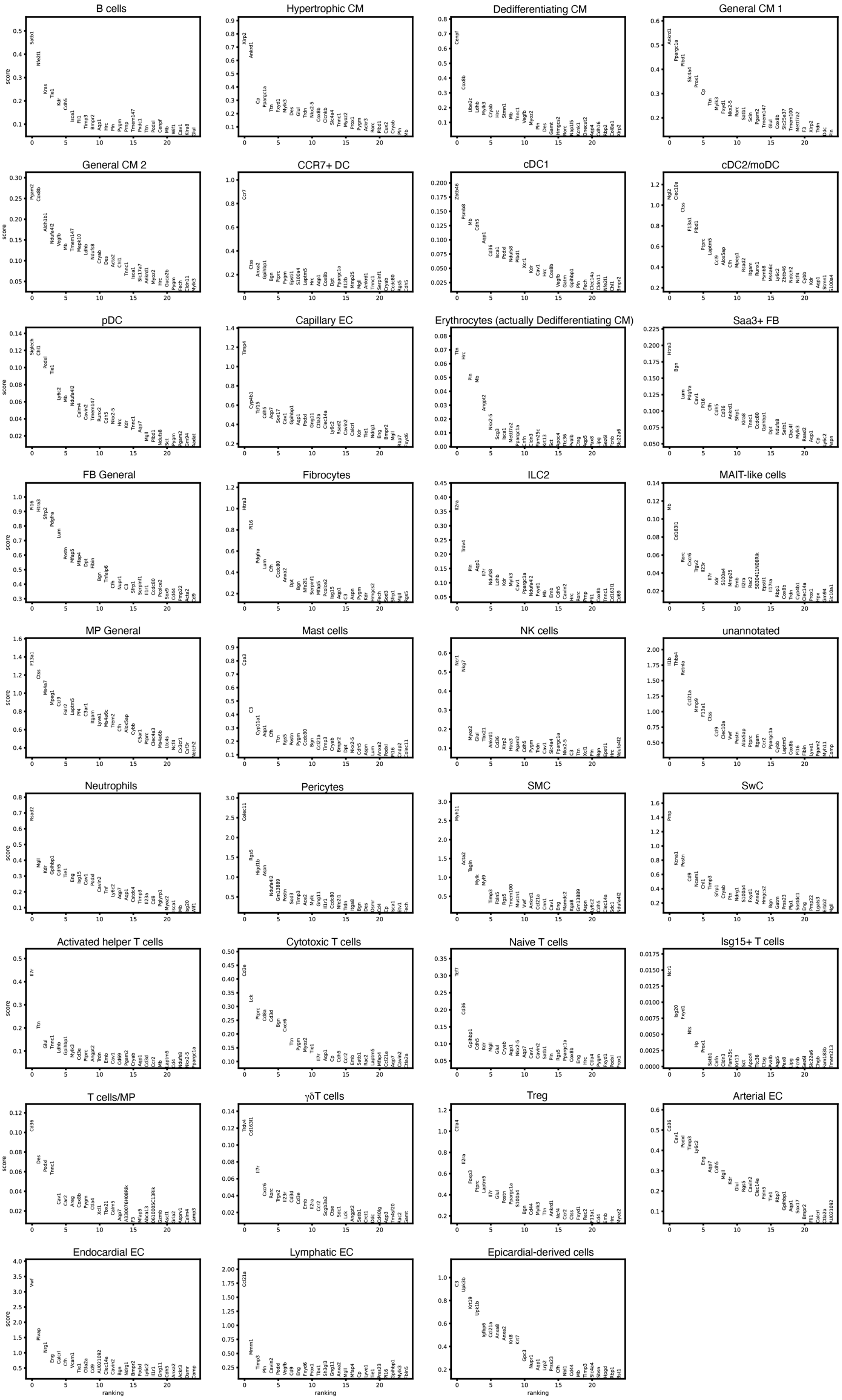
B cells
Hypertrophic CM
Dedifferentiating CM
General CM 1
General CM 2
CCR7+ DC
cDC1
cDC2/moDC
pDC
Capillary EC
Erythrocytes (actually Dedifferentiating CM)
Saa3+ FB
FB General
Fibrocytes
ILC2
MAIT-like cells
MP General
Mast cells
NK cells
unannotated
Neutrophils
Pericytes
SMC
SwC
Activated helper T cells
Cytotoxic T cells
Naive T cells
Isg15+ T cells
T cells/MP
γδT cells
Treg
Arterial EC
Endocardial EC
Lymphatic EC
Epicardial-derived cells
score
ranking

**Figure S2. Gene expression of spatial-annotated cell types**

Top 25 differentially expressed genes (DEGs) of NiCo-annotated cell types for the Cryo day 7 spatial data.

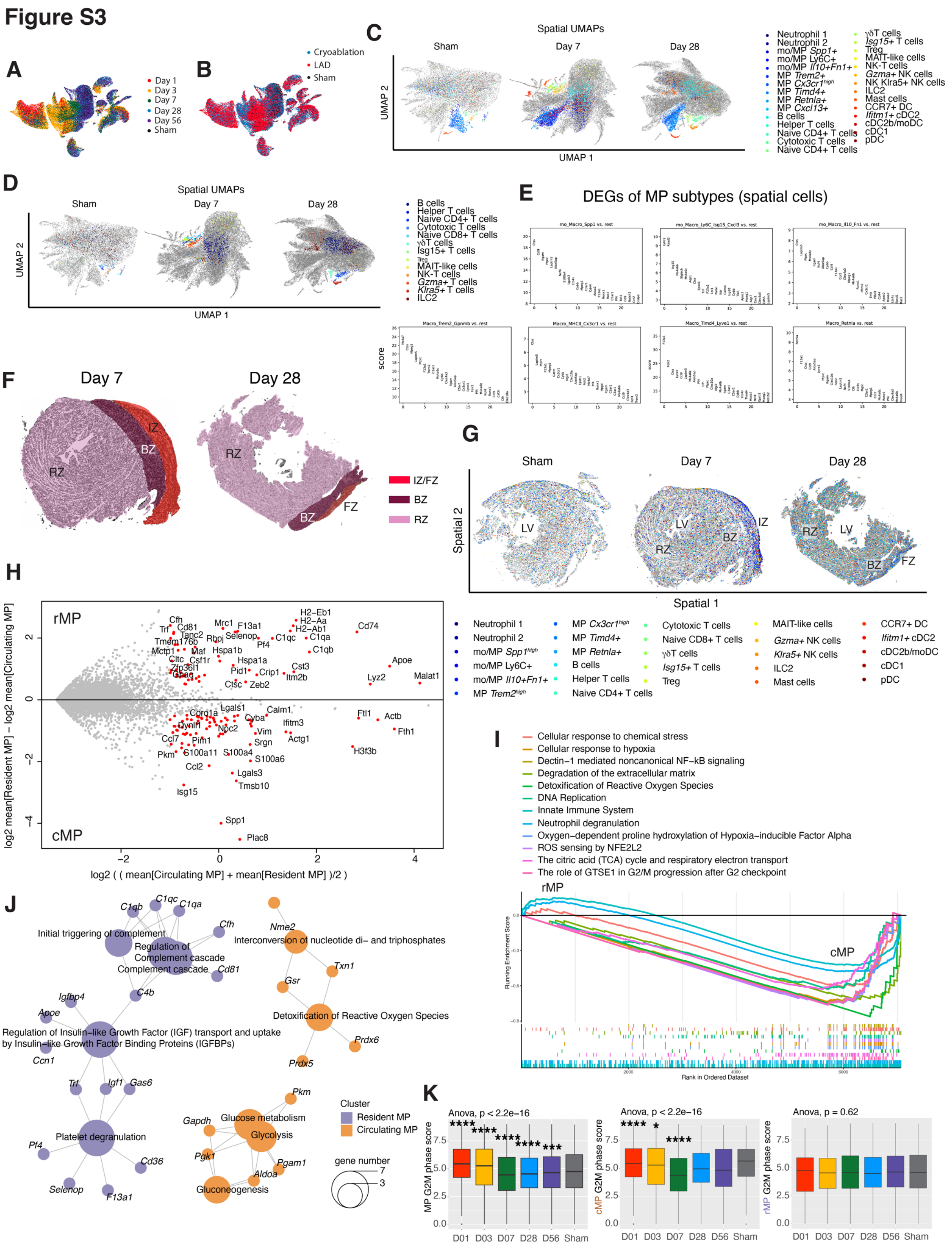


**Figure S3. Comparison of cMP and rMP gene expression profiles**

**A and B**, UMAP of immune cell populations grouped by time point post-lesion (**A**) or surgery types (**B**). **C and D**, spatial UMAP showing all NiCo-annotated immune cell (sub-)populations (**c**) or lymphocytes only (**D**). **E**, Top 25 DEGs of NiCo-annotated mo/MP cell types in the spatial data. **F**, definition of tissue regions for day 7 and day 28 samples, that were used for spatial cell type proportion analysis. **G**, spatial maps of all immune (sub-)populations in Sham, day 7, and day 28, under the detailed annotation scheme. **H and I**, differential gene expression analysis ($p < 0.05$, highlighted in red) (**H**) and gene set enrichment analysis (from ReactomePA database, min Gene Size = 15, max Gene-Size = 800, $p < 0.05$) (**I**) between rMP and cMP. **J**, gene concept network comparing differential pathways between rMP and cMP. Pathways are located at the center node of each cluster, while the corresponding genes are shown as connected smaller nodes. Sizes of pathway nodes correspond to the number of contributing genes (from ReactomePA database, differential gene expression log2-foldchange > 1). **K**, summed expressions of G2M cell cycle genes in all MP (top), cMP (middle) and rMP (bottom) across all time points post-lesion, and Sham. One-way ANOVA for multiple comparisons was performed. *$p < 0.05$, ** $p < 0.01$, *** $p < 0.001$, and **** $p < 0.0001$; ns, not significant.

#### Figure S4

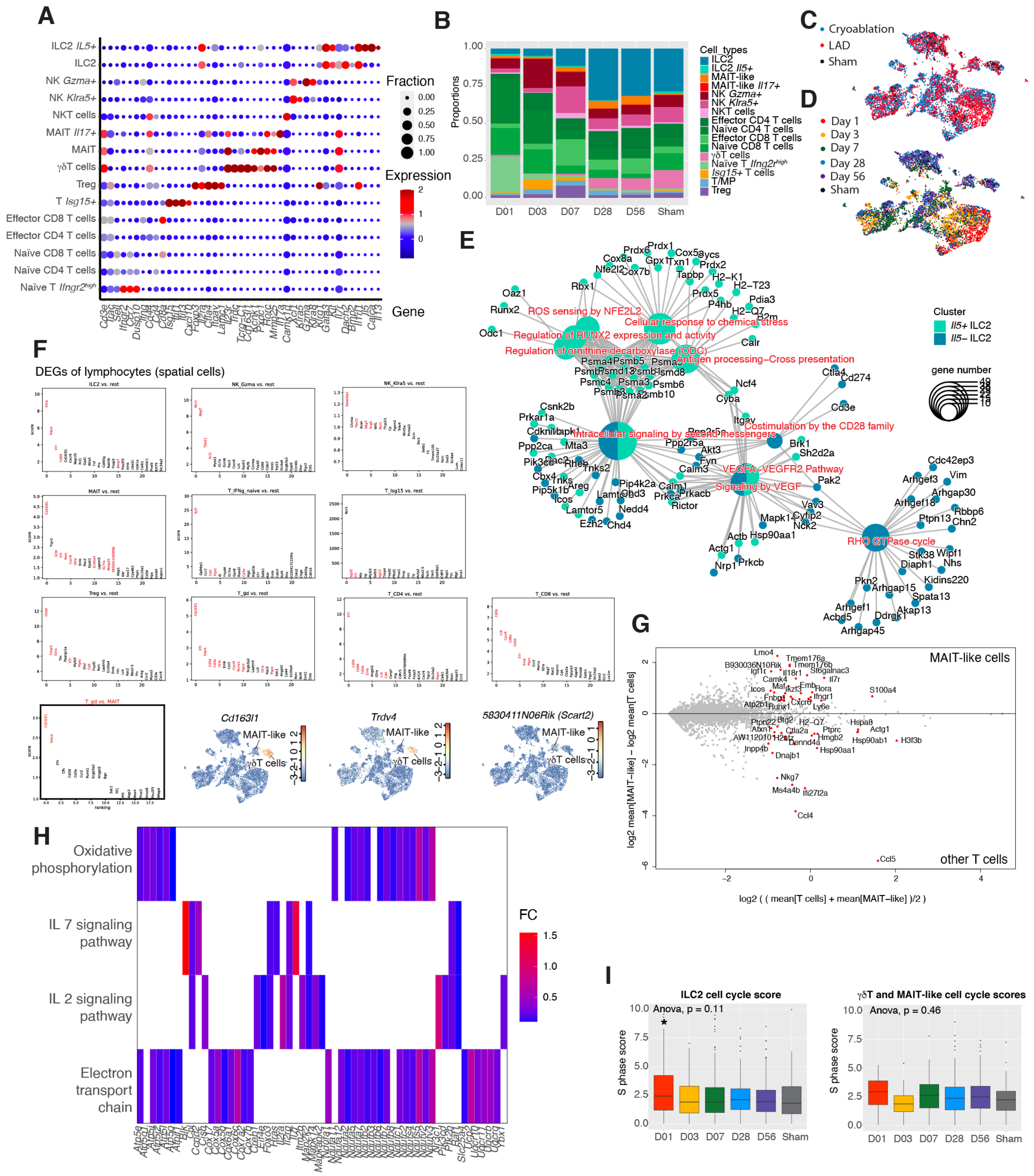


**Figure S4. Gene expression and pathway analysis in lymphocyte populations**

**A – D**, scRNA-seq of lymphocyte populations (without B cells) for LAD, cryoablation and Sham on days 1 to 56 post lesion. **A**, dot plot showing marker genes for each population. Dot diameter represents the fraction of cells of that cell type expressing the plotted gene. **B,** barplot showing cell type proportion dynamics of lymphocyte populations across days 1–56 and Sham. **C and D**, UMAPs of lymphocyte populations grouped by surgery types (**C**) and time points post-lesion (**D**). **E**, gene concept network comparing differential pathways between *Il5+* and *Il5-* ILC2. Pathways are located at the center node of each cluster, while the corresponding genes are shown as connected smaller nodes. Sizes of pathway nodes correspond to the number of contributing genes (from ReactomePA database, differential gene expression log2-

foldchange > 0.3). **F**, Top 25 DEGs of NiCo-annotated lymphocyte cell types in the spatial data. Text in red highlights known marker genes for the corresponding cell types. Lower panels, DEG plot and gene expression UMAPs of lymphocytes showing genes that are used to differentiate between $\gamma\delta$T cells and MAIT-like cells in the spatial data. **G**, differential gene expression analysis ($p < 0.05$, highlighted in red) between MAIT-like cells and other T cells. **H**, Heatmap plot of pathway enrichment analysis and corresponding genes enriched in MAIT-like cells against other T cells (ReactomePA database, DEG log2-foldchange > 0, $p < 0.05$). **I**, summed expressions of cell cycle genes (S phase) among ILC2, and $\gamma\delta$T cells + MAIT-like cells, across all time points post lesion, and Sham.

#### Figure S5

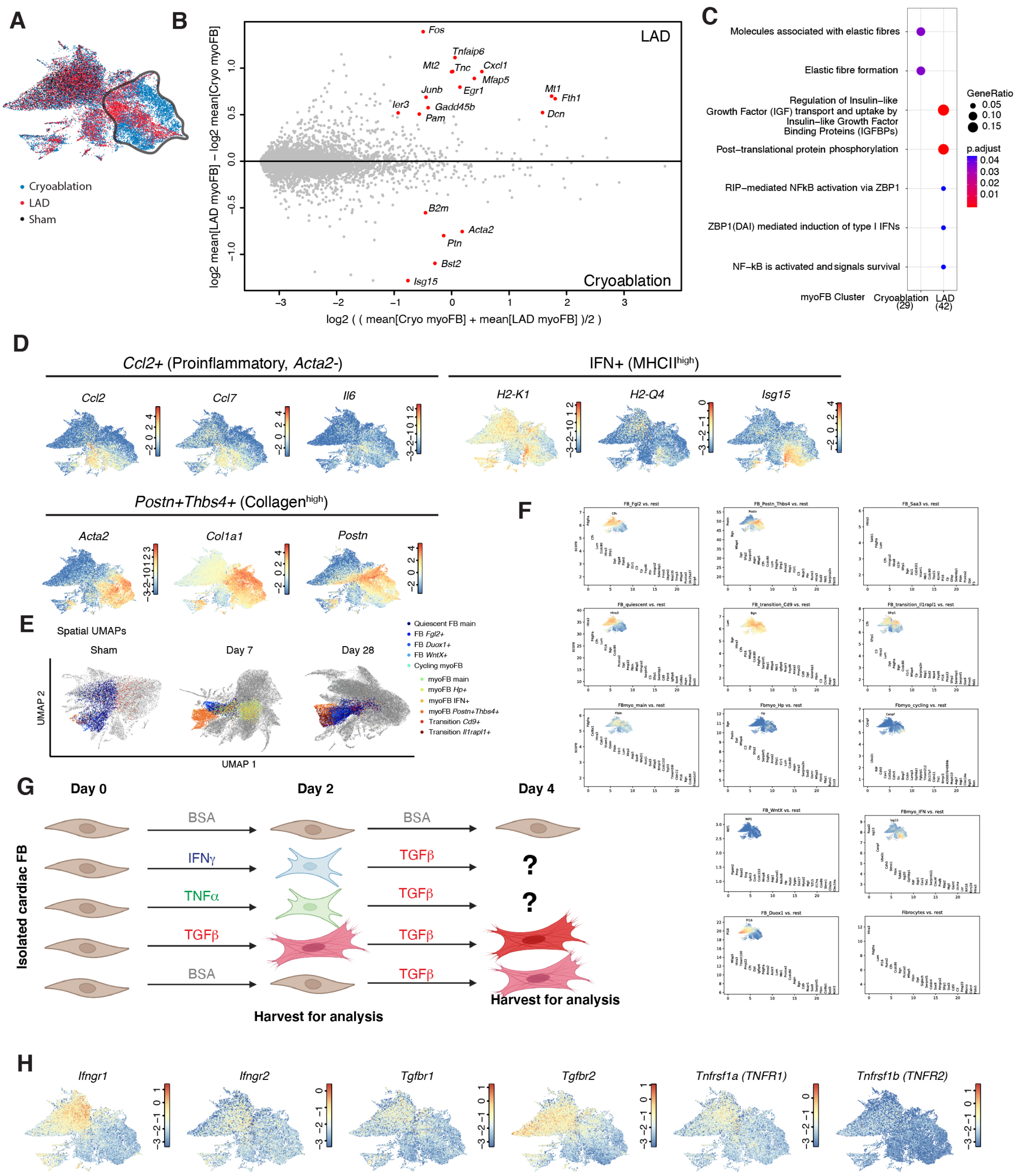

**Figure S5. Gene expression profiles of FB activation**

**A**, UMAP of FB populations grouped by surgery type. Highlighted cells are myoFB, where differential gene expression analysis ($p < 0.05$, indicated in red) between LAD and Cryoablation (**B**), as well as pathway enrichment analysis (from ReactomePA database, differential gene expression log2-foldchange > 0.4) comparing between the two groups (**C**) were performed. **D**, FB UMAP plotting marker genes for early proinflammatory *Ccl2+* FB, IFN+ MHCII$^{high}$ FB and *Postn+Thbs4+*Collagen$^{high}$ myoFB. **E**, spatial UMAP showing all NiCo-annotated FB subtypes, with detailed annotation scheme. **F**, top 25 DEGs of NiCo-annotated FB subtypes in the spatial data. In each plot, expression of one or more DEG(s) are also plotted in the corresponding scRNA-seq UMAP, showing enrichments of the same DEGs in the corresponding cell subtype across scRNA-seq and spatial data, indicating confident annotation of these cell subtypes. Alternatively, plots without any scRNA-seq UMAP indicate that the DEGs for this annotation do not match the corresponding cell subtype in the scRNA-seq data, and thus these annotations are not used for downstream analyses. **G**, schematic diagram of the isolated cardiac FB culturing with different stimulation factors in serum-free medium, TNF$\alpha$, IFN$\gamma$ and TGF$\beta$, respectively, for 2 days, followed by 2 more days of exposure to TGF$\beta$. Cells were collected at day 2 and 4 for qRT-PCR quantification to determine phenotypic changes. **H**, UMAP of FB populations showing expression of the receptor genes for the ligands tested in (**G**).

#### Figure S6

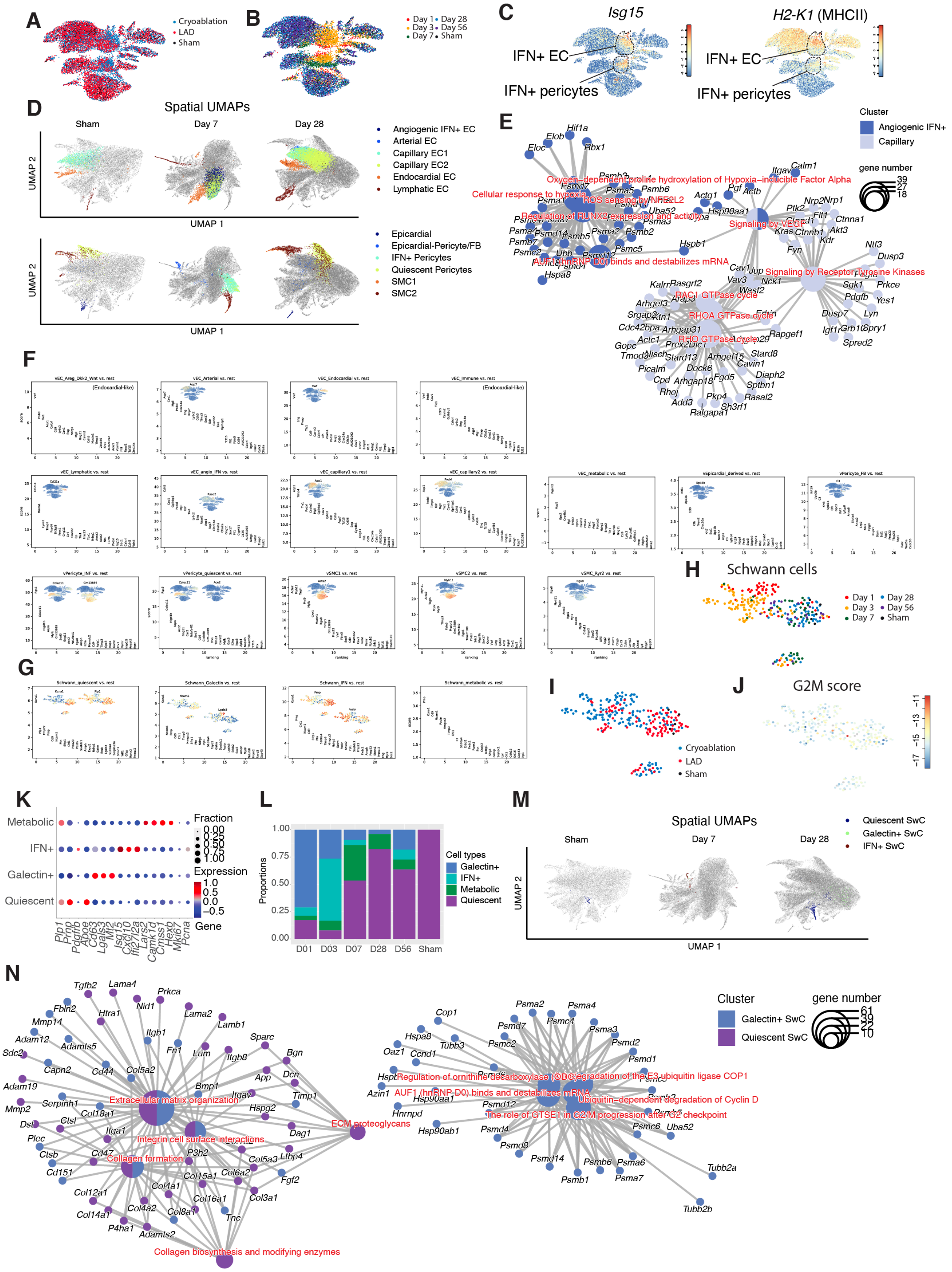


**Figure S6. Gene expression profiles and pathways of vascular cell and SwC populations**

**A and B**, UMAP of vascular populations were grouped by surgery types (**A**) or time points post-lesion (**B**). **C**, vascular cell UMAP plotting expression of IFN pathway gene *Isg15* and MHCII gene *H2-K1*. **D**, spatial UMAP showing all NiCo-annotated vascular cell types, with detailed annotation scheme. **E**, gene concept network comparing differential pathways between angiogenic IFN EC and capillary EC. Pathways are located at the center node of each cluster, while the corresponding genes are shown as connected smaller nodes. Sizes of pathway nodes correspond to the number of contributing genes (from ReactomePA database, differential gene expression log2-foldchange > 0.5). **F - G**, top 25 DEGs of NiCo-annotated vascular cell subtypes (**F**) and SwC subtypes (**G**) in the spatial data. In each plot, expression of one or more DEG(s) are also plotted in the corresponding scRNA-seq UMAPs, showing enrichments of the same DEGs in the corresponding cell subtype across scRNA-seq and spatial data, indicating confident annotations of these cell subtypes. Alternatively, plots without any scRNA-seq UMAP indicate that the DEGs for this annotation do not match the corresponding cell subtype in the scRNA-seq data, and thus these annotations are not used for downstream analyses. **H and I**, UMAP of SwC subtypes grouped by surgery types (**H**) or time points post-lesion (**I**). **J**, G2M score of SwC subtypes. **K**, dot plot showing DEGs in each SwC subtypes. **L**, barplot showing cell type proportion dynamics of SwC subtypes across days 1–56 and ham. **M**, spatial UMAP showing all NiCo-annotated SwC subtypes, with detailed annotation scheme. **N**, gene concept network comparing differential pathways between Galectin+ SwC and quiescent SwC. Pathways are located at the center node of each cluster, while the corresponding genes are shown as connected smaller nodes. Sizes of pathway nodes correspond to the number of contributing genes.

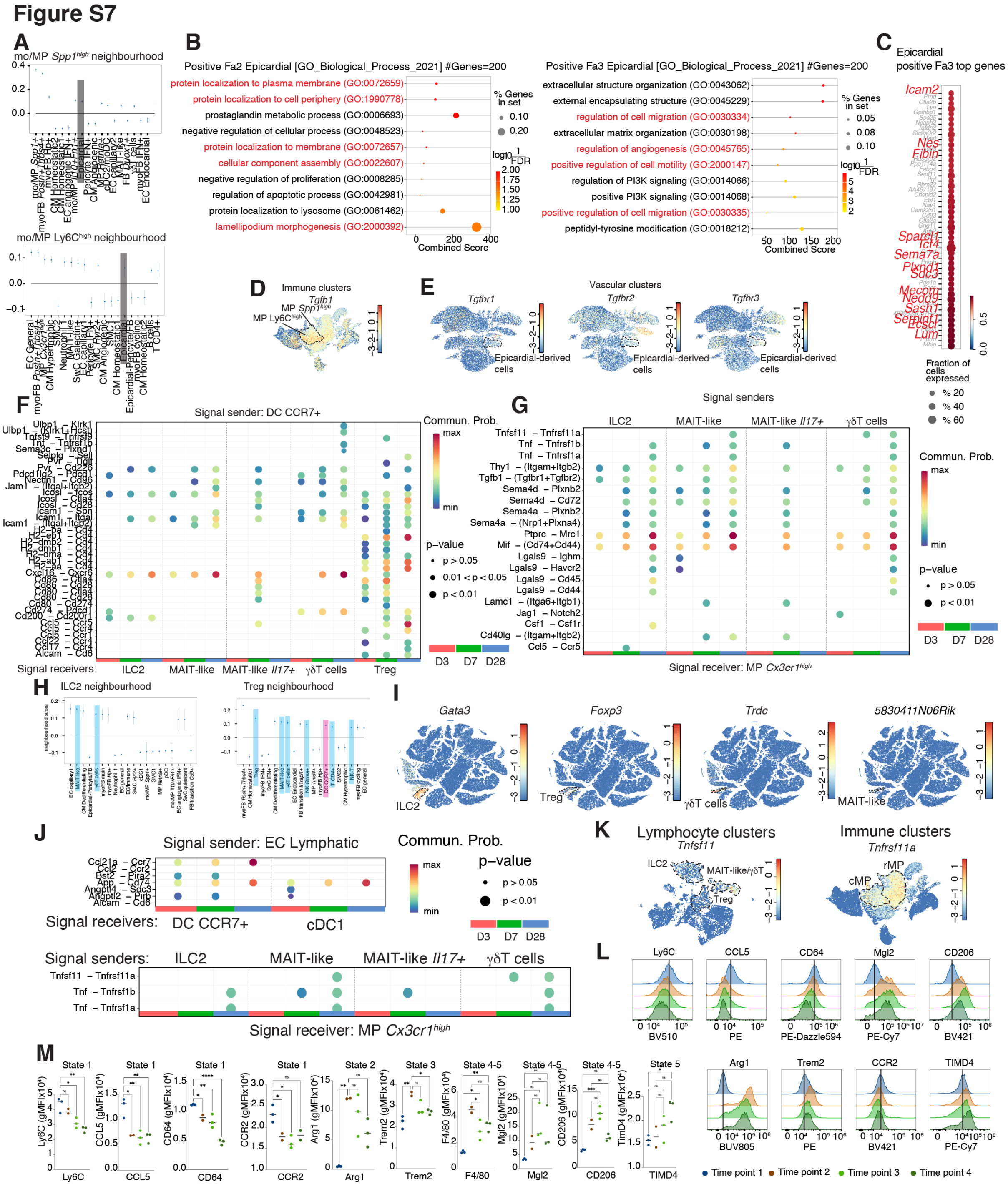

**Figure S7. Spatio-temporal cellular interactions in immune cell neighborhoods**

**A**, day 7 spatial neighborhoods of Ly6C$^{high}$ and *Spp1$^{high}$* mo/MP respectively. Highlighted in dark are epicardial cells. **B**, pathway enrichment analysis showing top pathways that are positively correlating with epicardial cell factor (Fa) 2 and 3, respectively. **C**, top genes that are positively correlated to epicardial cell Fa3. Genes and pathways highlighted in red are associated to cell migration and/or EMT. **D**, immune cell UMAP highlighting expression of *Tgfb1*. cMP are indicated. **E**, vascular cell UMAP highlighting TGFβ receptor genes. Epicardial-derived cells are indicated. **F – G**, CellChat ligand-receptor interaction predictions of CCR7+ DC to unconventional lymphocytes (**F**), and unconventional lymphocytes to *Cx3cr1$^{high}$* MP (**G**), from Day 3 – 28. **H**, Day 7 cell neighborhoods surrounding ILC2 and Treg, respectively. Cell types highlighted in red and blue are myeloid and lymphoid cells, respectively. **I**, UMAP of all cells showing markers for ILC2 (*Gata3*), Treg (*Foxp3*), $\gamma\delta$T cells (*Trdc*) and MAIT-like cells (*5830411N06Rik*) to indicate their location in the UMAP. **J**, CellChat ligand-receptor interaction predictions across Lymphatic EC, DC, unconventional T cells and $Cx3cr1^{high}$ MP, from days 3–28. **K**, UMAPs of lymphocyte (left) and all immune cell (right) populations showing expression of *Tnfsf11* (ligand) and *Tnfrsf11a* (receptor), respectively. Scale bar: log2-normalized gene expression. **L**, representative FACS histograms plotting the area-normalized expression of different MP cell state-associated surface proteins, across 4 culturing time points. Time point legend colors are indicated in (**M**). **M**, quantifications of cultured MP surface protein expression levels by Geometric mean fluorescence intensity (gMFI) from the histogram plots in represented in (**L**). Statistical significance was calculated by unpaired two-tailed t-test for two experimental groups. * $p < 0.05$, ** $p < 0.01$, *** $p < 0.001$, and **** $p < 0.0001$; ns, not significant.

#### Figure S8

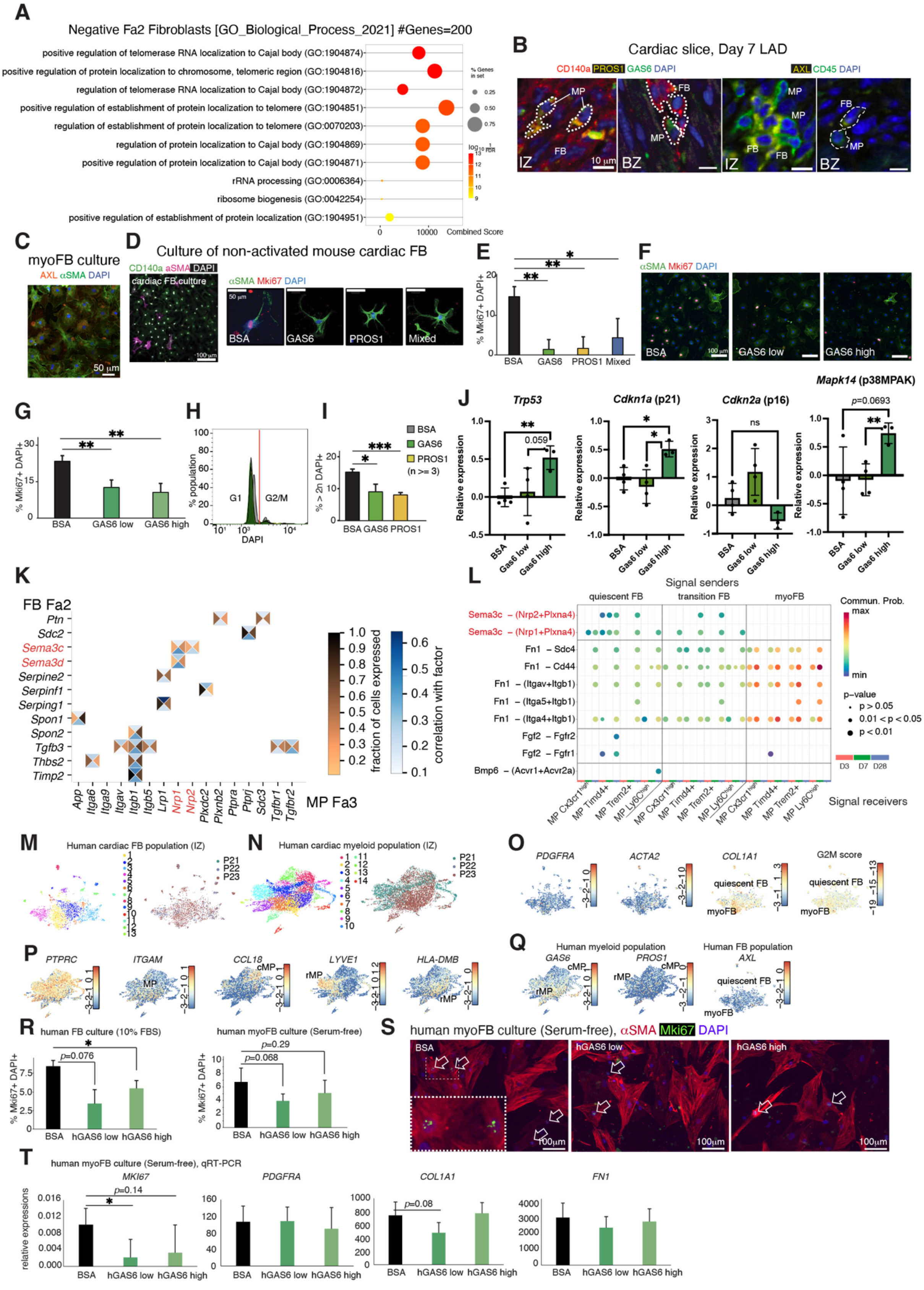

A
Negative Fa2 Fibroblasts [GO_Biological_Process_2021] #Genes=200
positive regulation of telomerase RNA localization to Cajal body (GO:1904874)
positive regulation of protein localization to chromosome, telomeric region (GO:1904816)
regulation of telomerase RNA localization to Cajal body (GO:1904872)
positive regulation of establishment of protein localization to telomere (GO:1904851)
regulation of establishment of protein localization to telomere (GO:0070203)
regulation of protein localization to Cajal body (GO:1904869)
positive regulation of protein localization to Cajal body (GO:1904871)
rRNA processing (GO:0006364)
ribosome biogenesis (GO:0042254)
positive regulation of establishment of protein localization (GO:1904951)
Combined Score
B
Cardiac slice, Day 7 LAD
CD140a PROS1 GAS6 DAPI
AXL CD45 DAPI
C
myoFB culture
AXL αSMA DAPI
D
Culture of non-activated mouse cardiac FB
CD140a aSMA DAPI
αSMA Mki67 DAPI
BSA GAS6 PROS1 Mixed
E
% Mki67+ DAPI+
F
αSMA Mki67 DAPI
BSA GAS6 low GAS6 high
G
% Mki67+ DAPI+
BSA GAS6 low GAS6 high
H
% population
G1 G2/M
DAPI
I
% > 2n DAPI+
BSA GAS6 PROS1 (n >= 3)
J
Trp53
Cdkn1a (p21)
Cdkn2a (p16)
Mapk14 (p38MPAK)
Relative expression
BSA Gas6 low Gas6 high
K
FB Fa2
Ptn Sdc2 Sema3c Sema3d Serpine2 Serpinf1 Serping1 Spon1 Spon2 Tgfb3 Thbs2 Timp2
App Itga6 Itga9 Itgav Itgb1 Itgb5 Lrp1 Nrp1 Nrp2 Plxdc2 Plxnb2 Ptpra Ptprj Sdc3 Tgfbr1 Tgfbr2
MP Fa3
fraction of cells expressed
correlation with factor
L
Signal senders
quiescent FB transition FB myoFB
Sema3c – (Nrp2+Plxna4)
Sema3c – (Nrp1+Plxna4)
Fn1 – Sdc4
Fn1 – Cd44
Fn1 – (Itgav+Itgb1)
Fn1 – (Itga5+Itgb1)
Fn1 – (Itga4+Itgb1)
Fgf2 – Fgfr2
Fgf2 – Fgfr1
Bmp6 – (Acvr1+Acvr2a)
Commun. Prob.
p-value
D3 D7 D28
Signal receivers
M
Human cardiac FB population (IZ)
N
Human cardiac myeloid population (IZ)
O
PDGFRA ACTA2 COL1A1 G2M score
quiescent FB myoFB
P
PTPRC ITGAM CCL18 LYVE1 HLA-DMB
Q
Human myeloid population
GAS6 PROS1
Human FB population
AXL
R
human FB culture (10% FBS)
human myoFB culture (Serum-free)
% Mki67+ DAPI+
BSA hGAS6 low hGAS6 high
S
human myoFB culture (Serum-free), αSMA Mki67 DAPI
BSA hGAS6 low hGAS6 high
T
human myoFB culture (Serum-free), qRT-PCR
MKI67 PDGFRA COL1A1 FN1
relative expressions
BSA hGAS6 low hGAS6 high

**Figure S8. MP-FB interactions in mouse and human hearts**

**A**, pathways negatively associated with FB Fa2 identified from the GO Biological Process 2021 database. **B**, IF staining of day 7 post-LAD IZ and BZ of cardiac slices, respectively. Dotted lines in left panels highlight MP (CD140a- PROS1+ GAS6+) and FB (CD140a+). Dotted lines in the right panels highlight cells resembling MP (CD45+ AXL+) and FB (CD45- AXL+). **C**, immunofluorescence staining of AXL in myoFB (αSMA+) cell culture. **D – I**, cardiac FB (non-TGFβ-activated) culture experiments. Mouse cardiac FB were pre-cultured in medium with 10% FBS overnight, then exposed in recombinant mouse GAS6, PROS1, both (mixed), or BSA in serum-free medium for 48 hours. Cells were then harvested for immunofluorescence staining with CD140a, aSMA and Mki67 (**D**), followed by quantification of CD140a+ Mki67+ nuclei (**E**). FB were cultured in different concentrations of GAS6, immunofluorescence-stain (**F**) and quantified (**G**). **H** and **I**, FB cultured with GAS6 or PROS1 were harvested for FACS quantification of percentage of nuclei with > 2n DNA content, indicated by G1 and G2M peaks of DAPI intensity (**H**). **J**, qRT-PCR quantification of senescence and cell cycle arrest marker genes from cultured mouse isolated cardiac myoFB exposed to GAS6 ligand. **K**, Ligand-receptor interactions, where the ligand and receptor correlate to FB Fa2 and MP Fa3, respectively. Boxes depict correlation with factors (blue) and fraction of cells expressing the gene (brown) **L**, CellChat ligand-receptor interaction prediction from days 3–28. Signal senders are various FB subtypes, signal receivers are various MP subtypes. In **K** and **L**, Sema3-Nrp interactions are highlighted in red. **M–Q**, re-analysis of published snRNA-seq data from 3 MI patients (P21, 22 and 23), isolated from IZ (Kuppe et al, 2022). **M** and **N**, human cardiac FB (**M**) and myeloid (**N**) populations. Nuclei are grouped according to cluster numbers (left panels) and patient identities (right panels). **O–Q**, UMAP plots showing expressions of FB G2M score/marker genes (**O**), MP marker genes (**P**), and expression of *GAS6*, *PROS1* and *AXL* in these two populations (**Q**). Quiescent FB were defined by *PDGFRA+ ACTA2$^{low}$ COL1A1$^{low}$* and lower G2M score. myoFB were defined by *PDGFRA+ ACTA2$^{high}$ COL1A1$^{high}$* and higher G2M score. cMP were defined by *PTPRC+ ITGAM+ CCL18+ HLA-DMB$^{low}$ LYVE1-*, rMP were defined by *PTPRC+ ITGAM+ CCL18- HLA-DMB$^{high}$/LYVE1+*. **R–T**, purified human primary cardiac FB culture experiments. FB were either exposed to two doses of recombinant human GAS6 ligand or BSA, in serum-free medium, 10% FBS, or serum-free with cells pre-activated into myoFB by recombinant human TGFβ overnight. After 48 hours of GAS6 exposure, cells were harvested for immunofluorescence staining and qRT-PCR quantifications. **R**, percentages of proliferating cells (Mki67+ DAPI+). **S**, representative images of myoFB cultures, stained with αSMA and Mki67 antibodies. **T**, qRT-PCR quantifications of *MKI67, PDGFRA, COL1A1* and *FN1* for myoFB culture, quantified relative to reference gene *GAPDH*. For (E), (G), (I), (J), (R) and (T), statistical significance was calculated by unpaired two-tailed t-test for two experimental groups. * $p < 0.05$, ** $p < 0.01$, *** $p < 0.001$; ns, not significant.

#### Figure S9

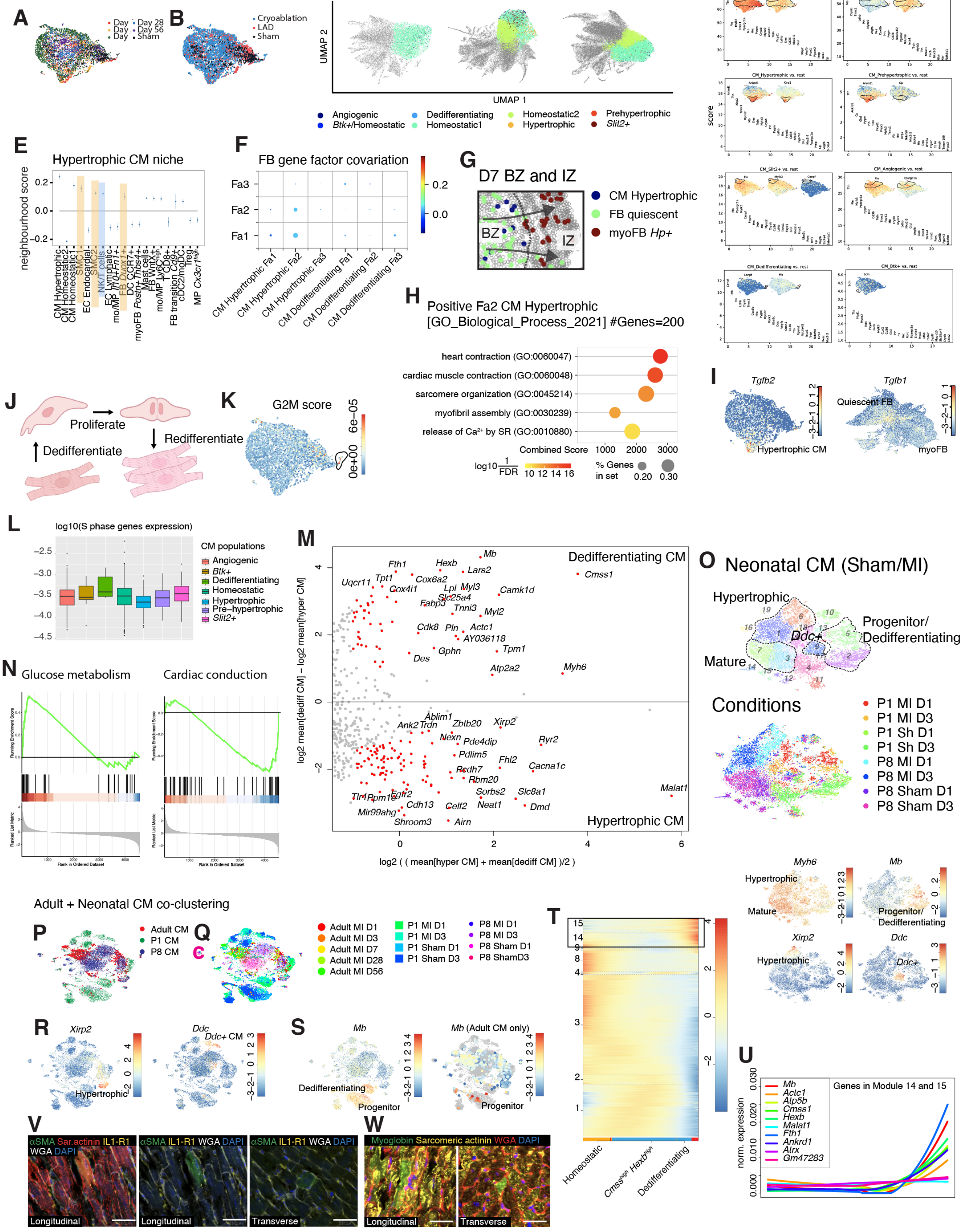


**Figure S9. Comparison of dedifferentiating and hypertrophic CM**

**A and B**, UMAP of CM populations grouped by time points post-lesion (**A**) or surgery types (**B**). **C**, spatial UMAP showing all NiCo-annotated CM subtypes. **D**, top 25 DEGs of NiCo-annotated CM subtypes in the spatial data. In each plot, expression of one or more DEG(s) are also plotted in the corresponding scRNA-seq UMAP(s), showing enrichments of the same DEGs in the same cell subtype across scRNA-seq and spatial data, indicating confident annotations of these cell subtypes. **E–I**, hypertrophic CM – FB interaction in Day 7 BZ. **E**, spatial neighborhood of hypertrophic CM on Day 7. **F**, spatial gene factor co-variation dot plot of FB neighborhood on Day 7 heart. Only interactions with CM subtypes are shown. Circle size scales linearly with $-\log_{10}$(p-value), and circle color indicates ridge regression coefficients. **G**, spatial map showing hypertrophic CM – FB neighbourhood in the BZ. Arrows indicate direction of FB migration and maturation. **H**, pathways positively associated with hypertrophic CM Fa2 identified by GO Biological Process 2021 database. **I**, CM (left) and FB (right) UMAPs plotting expressions of TGFβ ligands. **J**, schematic diagram showing dedifferentiation-proliferation-redifferentiation trajectory of CM upon MI. **K**, UMAP of CM population showing G2M scores. Circled cells are the dedifferentiating CM. **L**, quantification of log10-transformed summed S phase cell cycle gene expression across all CM subtypes. **M**, differential gene expression analysis and between dedifferentiating CM and hypertrophic CM. Significant genes are indicated in red ($p < 0.05$) **N**, gene set enrichment analysis comparing between dedifferentiating CM and hypertrophic CM. **O**, re-analysis of the published neonatal post-MI CM datasets from Cui et al. (2020). Top panel, UMAP clustering of CM subtypes. Middle panel, condition of CM samples. D1, day 1 post-MI/Sham. D3, day 3 post-MI/Sham. bottom panels, expression plots of gene markers for mature (*Myh6$^{high}$*), progenitor/dedifferentiating (*Mb$^{high}$*), hypertrophic (*Xirp2+*) and Ddc+ (*Ddc+*) CM subtypes. **P**, UMAP co-clustering of our adult CM dataset (labelled in red) with neonatal post-MI CM datasets. P1 hearts are regenerative (green) while P8 hearts are non-regenerative (blue). **Q**, UMAP showing detailed CM conditions. **R and S**, marker genes annotating hypertrophic (*Xirp2+*) and *Ddc+* CM, which do not contribute to heart regeneration (**R**), and progenitor and dedifferentiating CM (*Mb$^{high}$*), which are the main contributors of heart regeneration (**S**). Right panel in **S**, same UMAP where only adult CM are shown. Adult dedifferentiating CM (*Mb$^{high}$*) co-cluster with P1 CM progenitor cells. **T**, FateID pseudo-time expression profiles derived by self-organizing maps (SOM). Heatmap showing gene expression profiles which were grouped into 14 modules. Highlighted modules (14 and 15) are genes with gradually increasing expression across the dedifferentiation trajectory. **U**, pseudo-time expression profiles of example genes from modules 14 and 15. **V and W**, IF staining of day 7 LAD cardiac slices in the RZ. Dedifferentiating CM are identified by Sarcomeric actinin+ aSMA+ and IL1-R1+ (**V**) or Sarcomeric actinin+ Myoglobin+ (**W**). WGA labels cell membranes.

#### Figure S10

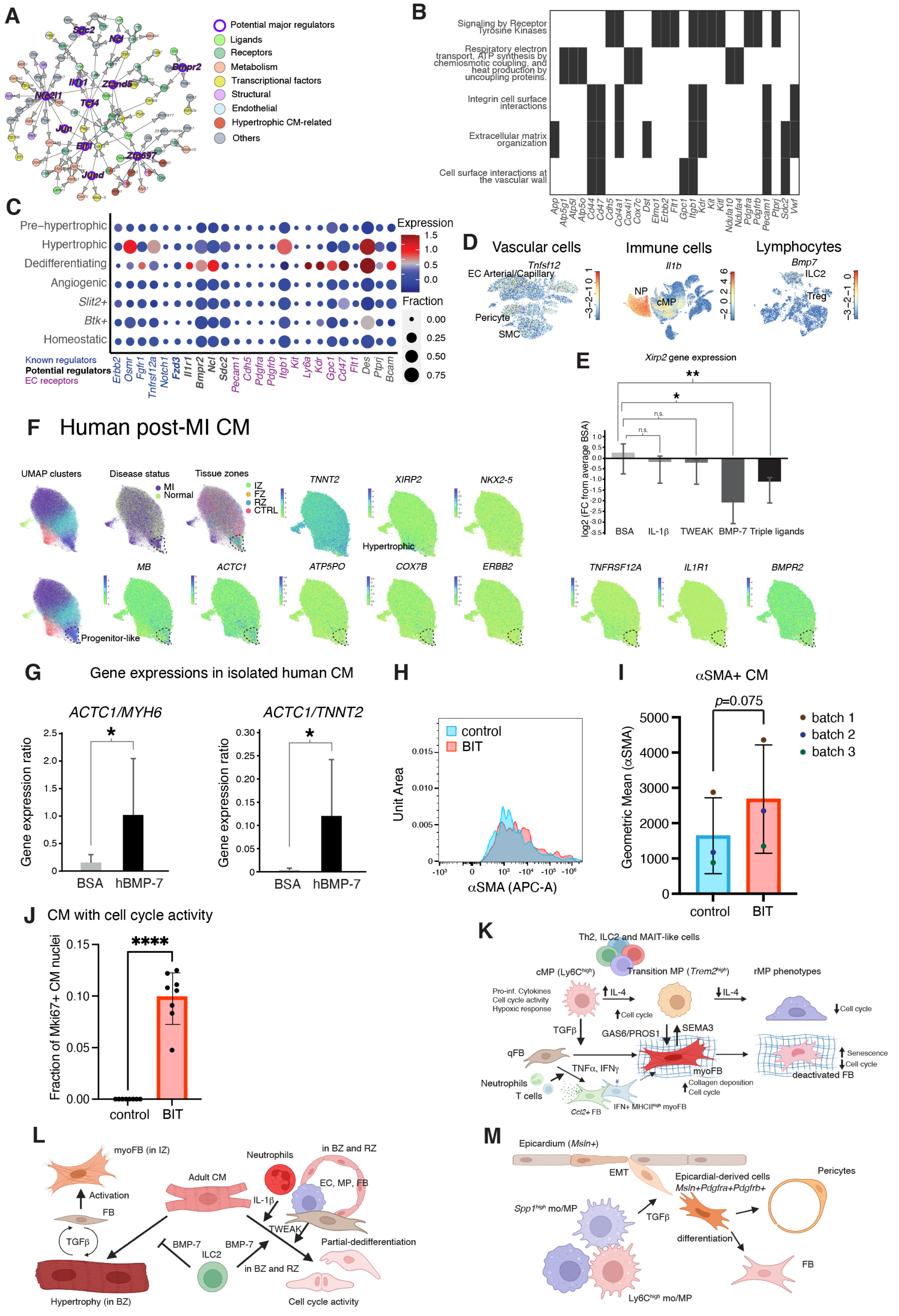

A
Potential major regulators
Ligands
Receptors
Metabolism
Transcriptional factors
Structural
Endothelial
Hypertrophic CM-related
Others
B
Signaling by Receptor Tyrosine Kinases
Respiratory electron transport, ATP synthesis by chemiosmotic coupling, and heat production by uncoupling proteins.
Integrin cell surface interactions
Extracellular matrix organization
Cell surface interactions at the vascular wall
C
Pre–hypertrophic
Hypertrophic
Dedifferentiating
Angiogenic
Slit2+
Btk+
Homeostatic
Expression
Fraction
Known regulators
Potential regulators
EC receptors
D
Vascular cells
Immune cells
Lymphocytes
E
Xirp2 gene expression
log2 (FC from average BSA)
BSA
IL-1β
TWEAK
BMP-7
Triple ligands
F Human post-MI CM
UMAP clusters
Disease status
Tissue zones
Hypertrophic
Progenitor-like
G
Gene expressions in isolated human CM
ACTC1/MYH6
ACTC1/TNNT2
Gene expression ratio
BSA
hBMP-7
H
control
BIT
Unit Area
αSMA (APC-A)
I
αSMA+ CM
p=0.075
batch 1
batch 2
batch 3
Geometric Mean (αSMA)
control
BIT
J
CM with cell cycle activity
Fraction of Mki67+ CM nuclei
control
BIT
K
Th2, ILC2 and MAIT-like cells
cMP (Ly6C^high)
Transition MP (Trem2^high)
rMP phenotypes
Pro-inf. Cytokines
Cell cycle activity
Hypoxic response
IL-4
Cell cycle
TGFβ
GAS6/PROS1
SEMA3
qFB
TNFα, IFNγ
myoFB
Neutrophils
T cells
Collagen deposition
Cell cycle
Ccl2+ FB
IFN+ MHCII^high myoFB
Senescence
deactivated FB
L
myoFB (in IZ)
Activation
FB
TGFβ
Hypertrophy (in BZ)
Adult CM
Neutrophils
in BZ and RZ
EC, MP, FB
IL-1β
TWEAK
BMP-7
ILC2
in BZ and RZ
Partial-dedifferentiation
Cell cycle activity
M
Epicardium (Msln+)
EMT
Epicardial-derived cells
Msln+Pdgfra+Pdgfrb+
Pericytes
Spp1^high mo/MP
TGFβ
differentiation
FB
Ly6C^high mo/MP

**Figure S10. Gene program and activation of mouse and human CM dedifferentiation**
**A**, GENIE3 gene-regulatory network in CM undergoing dedifferentiation, deduced from SOM module 14 and 15 genes. **B**, enriched pathways and corresponding genes in module 14 and 15 (ReactomePA database, $p$-value < 0.05). **C**, dot plot showing expression of candidate dedifferentiation-promoting receptor genes in CM subtypes. **D**, UMAPs showing expression of candidate dedifferentiation-promoting ligand genes in vascular cells (left), all immune cells (middle) and lymphocytes (right). **E**, qRT-PCR of *Xirp2* (hypertrophic CM marker gene) in cultured CM after ligand exposure. log2-transformed relative gene expression fold change against BSA control are plotted. Unpaired two-tailed t-test was performed in each group against BSA control, where * $p < 0.05$, ** $p < 0.01$. **F**, human post-MI CM UMAP from snRNA-seq data by Kuppe et al. (2022) deposited in CELLxGENE. Marker genes for general CM (*TNNT2*), hypertrophic CM (*XIRP2*), and dedifferentiating CM (*NKX2-5, MB, ACTC1, ATP5PO* and *COX7B*) are plotted. Receptor genes identified in the mouse CM dedifferentiation niche are also plotted (*TNFRSF12A, IL1R1* and *BMPR2*). **G**, qRT-PCR of *ACTC1* (CM progenitor marker gene), *MYH6* and *TNNT2* (mature CM marker genes) in primary human CM progenitor-like cells after hBMP-7 exposure. *ACTC1*-to-*MYH6* and *ACTC1*-to-*TNNT2* gene expression ratios (as a progenitor state score) were compared against BSA control. **H and I**, αSMA expression in human cardiac slice CM. Representative FACS histogram plot showing the intensities of αSMA in control (BSA-treated) and BIT-treated CM (sarcomeric actinin+) (**H**), and the quantifications of the signal intensities (**I**). **J**, quantification of CM cell cycle activities (Sarcomeric actinin+ cell with Mki67+ nuclei). For (G), (I) and (J), statistical significance was calculated by unpaired two-tailed t-test for two experimental groups. * $p < 0.05$, ** $p < 0.01$, *** $p < 0.001$, and **** $p < 0.0001$; ns, not significant. **K–M**, Summary diagrams of the identified spatial-temporal niches in cardiac wound healing. **K**, Time-dependent interactions of MP, lymphocytes and FB in the fibrotic niche. At days 1–7 post-lesion, cMP promote differentiation of quiescent FB (qFB) and *Ccl2+* FB into myoFB via TGFβ. TNFα and IFNγ from neutrophils and T cells, respectively, can drive qFB into proinflammatory *Ccl2+* and IFN+ MHCII$^{high}$ phenotypes, respectively. The *Ccl2+* state can be further converted into myoFB upon TGFβ exposure. On Day 7, cMP gradually acquire an rMP phenotype through a transitional *Trem2$^{high}$* state, through the transient exposure of IL4+ from lymphocytes like Th2, ILC2 and MAIT-like cells. These transition and rMPs express GAS6 and PROS1 that dampen cell cycle rate in myoFB. Reciprocally, myoFB expresses SEMA3 which dampens the cell cycle rate in rMP. Eventually, this contributes to the reduction of both MP and myoFB in FZ at the later stage of healing, preventing excessive fibrosis and prolonged tissue remodeling.

**L**, CM hypertrophic and dedifferentiation niches. In the BZ early after lesion, adult CM colocalize with migrating FB, that produce TGFβ1, inducing CM into hypertrophy. In return, hypertrophic CM produce TGFβ2, which promotes the differentiation of migrating FB into myoFB as they reach IZ. Alternatively, in rare cases when CM simultaneously interact with myeloid cells, ILC2 and EC/FB, which provide IL-1β, BMP-7 and TWEAK, respectively, this results in formation of the CM dedifferentiation niche. Individually, IL-1β and TWEAK each provide marginal pro-proliferative effects. BMP-7 prevents CM from acquiring a hypertrophic phenotype. The synergistic effect of these interactions promotes CM dedifferentiation and cell cycle re-entry.

**M**, MP - epicardial-derived cell interaction. In IZ on day 7, MP promote epicardial-derived cell EMT gene expression profiles via TGFβ signaling.
